# *Ce*Lab, a Microfluidic Platform for the Study of Life History Traits, reveals Metformin and SGK-1 regulation of Longevity and Reproductive Span

**DOI:** 10.1101/2023.01.09.523184

**Authors:** Salman Sohrabi, Vanessa Cota, Coleen T. Murphy

**Affiliations:** Department of Molecular Biology, Princeton University, Princeton NJ 08544; LSI Genomics, Princeton University, Princeton NJ 08544; Department of Bioengineering, The University of Texas at Arlington, Arlington TX 76010

## Abstract

The potential to carry out high-throughput assays in a whole organism in a small space is one of the benefits of *C. elegans*, but worm assays often require a large sample size with frequent physical manipulations, rendering them highly labor-intensive. Microfluidic assays have been designed with specific questions in mind, such as analysis of behavior, embryonic development, lifespan, and motility. While these devices have many advantages, current technologies to automate worm experiments have several limitations that prevent widespread adoption, and most do not allow analyses of reproduction-linked traits. We developed a miniature *C. elegans* lab-on-a-chip device, *Ce*Lab, a reusable, multi-layer device with 200 separate incubation arenas that allows progeny removal, to automate a variety of worm assays on both individual and population levels. *Ce*Lab enables high-throughput simultaneous analysis of lifespan, reproductive span, and progeny production, refuting assumptions about the Disposable Soma hypothesis. Because *Ce*Lab chambers require small volumes, the chip is ideal for drug screens; we found that drugs previously shown to increase lifespan also increase reproductive span, and we discovered that low-dose metformin increases both. *Ce*Lab reduces the limitations of escaping and matricide that typically limit plate assays, revealing that feeding with heat-killed bacteria greatly extends lifespan and reproductive span of mated animals. *Ce*Lab allows tracking of life history traits of individuals, which revealed that the nutrient-sensing mTOR pathway mutant, *sgk-1*, reproduces nearly until its death. These findings would not have been possible to make in standard plate assays, in low-throughput assays, or in normal population assays.

## Introduction

*C. elegans* is a useful model system to study a variety of biological characteristics, but it is particularly well-suited to the study of aging because of its short lifespan and obvious, visible changes with age. Its genetic tractability, short generation time (<3 days), large progeny number (300 if self-fertilized, >600 if mated) and high degree of evolutionary conservation has allowed the discovery of conserved genetic regulators of development, longevity, reproductive aging, health span, and other important life history phenomena. Its small size (<1mm) and simple laboratory diet (*E. coli*) enables the maintenance of large numbers of animals in a small area. Together, these properties make *C. elegans* ideal for high-throughput experimentation and discovery applications, particularly for life history traits such as lifespan, progeny production, and reproductive span.

*C. elegans* assays often require a large sample size and frequent physical manipulations, rendering them highly labor-intensive^1^. Some of these challenges have been met by long-term culturing and imaging^2,3^ that have been used for large-scale lifespan analyses^4–6^. For example, a recent high-throughput analysis of lifespan and motility traits^7^ re-emphasized that induced Maximum Velocity best reflects health of individuals with age, as we previously reported^8^. However, such large-scale approaches usually require the elimination of progeny, limiting the analysis to lifespan and healthspan metrics that ignore reproductive traits and reproductive effects on lifespan. Therefore, other approaches are necessary to assess healthspan parameters associated with reproduction, such as progeny production, reproductive span, and post-mating lifespan^9,10^. Preparing growth medium in well plates for large-scale cultivation can potentially be a viable alternative, as it offers flexibility in experimental design for users to simultaneously test various biological conditions. However, in smaller wells, worms are easily lost to desiccation on the walls. Moreover, users are still required to carry out laborious transfer of worms. To address these problems, liquid assays in well-plates can expedite daily handling. However, worms in liquid alternate between stressful swimming and long quiescence states^11,12^ and they may also experience hypoxic conditions that compromise long-term incubation.

Microfluidic technologies have emerged as state-of-the-art tools to expedite worm assays, which can also offer a higher degree of control and accuracy. The integration of novel lab-on-chip devices with programmable valves, motorized platforms, image-based screening Sohrabi & Cota and Murphy, 2023– preprint version –www.biorxiv.org methods, and computer-assisted analysis has enabled faster development of new disease models and rapid phenotypic screening of large drug libraries^13^. Pioneering microfluidic chips have been designed for worm culture^14^, embryonic developmental studies^15,16^, microsurgery^17^, behavioral screening^18^, and high-content imaging^19,20^. Although advanced, these technologies are often assay-specific and single-use, and have limited capacity and flexibility in experimental design (Table S1). Moreover, they are often complicated to operate and slower than traditional assays, as new devices or chips are required for each experiment. Despite excellent proof-of-concept work, these drawbacks have prevented the widespread adoption of this technology.

Here we describe our development of *Ce*Lab, a microfluidic device for the study of *C. elegans* life history traits. Because *Ce*Lab solves problems of previous devices and plates, we were able to use it to discover new aspects of biology, as well. To automate multiple manual assays using one device, we have developed a reusable device that combines features of multi-well plates, microfluidic chips, and manual assays to address the limitations of previous technologies while retaining their best qualities. *Ce*Lab is a plug-and-play device that uses novel techniques for worm loading and operation of the device to close the gap between engineering design of microstructures and macro-world of conventional worm assays. Our device is highly reliable, versatile, and easy to use, with minimal operational costs and environmental impact, and offers high flexibility in experimental design. With daily bacterial feeding and daily manual scoring on a benchtop microscope, it closely resembles plate assays in its output, but offers individual worm information that is not possible in population plate assays. We found that carrying out lifespan and reproductive span assays using *Ce*Lab is ∼6x faster than corresponding plate assays (Table 1) which can significantly expedite worm research, enabling new biological discoveries. *Ce*Lab allowed us to discover that individual animals are not subject to the Disposable Soma hypothesis of resource utilization, and that heat-killed bacteria extend lifespan and reproductive span through specific genetic mechanisms. Finally, we found that mutation of *sgk-1*, the Serum- and Glucocorticoid-inducible kinase homolog, greatly extends reproductive span and reverses lifespan shortening normally observed after mating. These biological observations were possible because *Ce*Lab enables high-throughput population and individual analyses of multiple healthspan characteristics in reproductive animals simultaneously.

**Table 1:**
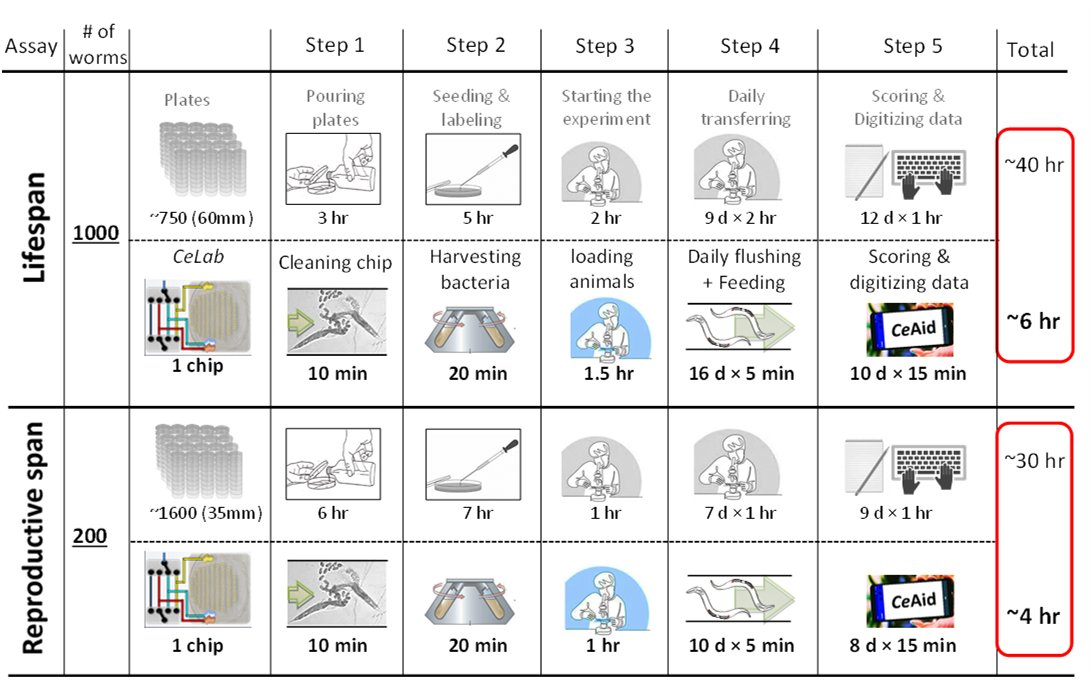
*Ce*Lab accelerates experimental discovery.

## Results

We are interested in the high-throughput measurement of lifespan, reproductive span, progeny production, and health metrics of individual animals, all of which requires long-term automated monitoring of individuals, daily removal of progeny, easy loading of many animals, and visualization of health span characteristics that are comparable to those seen on plates with manual manipulation. To meet these requirements, we needed to develop a new device.

### Design and use of *Ce*Lab

We designed and fabricated a multilayered Polydimethylsiloxane (PDMS) device for long-term *C. elegans* incubation and monitoring (Figure 1a-g, Figure S1a-d). One of the first requirements for long-term maintenance of *C. elegans* in microfluidic chambers is the presence of pillars^21^ (Figure 1a), as thrashing worms in pillarless environments are stressed and have shorter lifespans^14^. Hexagonally-arranged microposts allow animals to move in the same fashion as they do on an agar plate^18^ (Figure 1a). Imaging of the worms inside the chamber (Figure 1d) enables measurements of size and health span.

**Figure 1:**
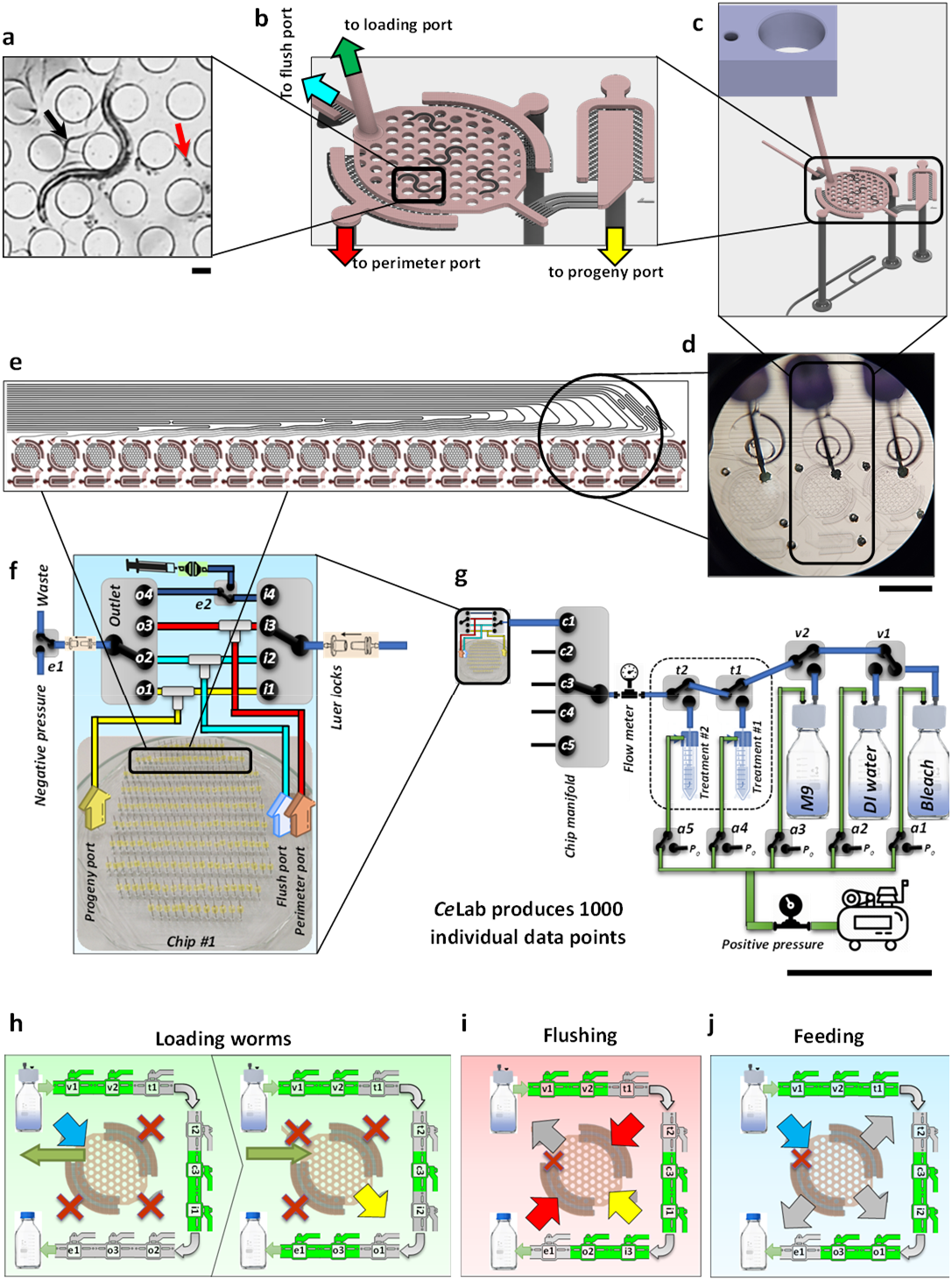
*Ce*Lab, a miniaturized *C. elegans* Lab on a chip. (a) Microposts, 200 µm in diameter with 300 µm spacing, allow animals crawl at a similar speed, wavelength, and amplitude as they do on an agar plate. Red and black arrows show fertilized egg and hatched progeny, respectively. Scale bar, 100 µm. (b) Each 3 mm diameter circular chamber can house multiple adult worms. Hatched progeny can be trapped in the “progeny chamber” to quantify daily progeny production. Unit number can be found next to the progeny chamber. (c) A 3D model of an incubation unit showing how different layers are connected. (d) Top-view image of wells and incubation area where pins block the loading ports. The smaller well, 1.5 mm in diameter, inside the 3 mm diameter main well, facilitates the worm loading process. Scale bar, 1 mm. (e) Identical hydraulic resistance across parallel channels connecting incubation arenas to flush, perimeter, and progeny ports ensures uniform flow distribution. (f) Flush, Perimeter, and Progeny ports can be either inlet, outlet, or closed by manipulation of on-chip inlet and outlet manifolds. Manipulating valves of inlet and outlet manifolds can generate various flow patterns inside the chip. (g) The main feeding line valve system of the *Ce*Lab control center directs flow from one reservoir into one chip at a time. Scale bar, 1 mm. (h-j) The schematics of the *Ce*Lab fluidic workflow demonstrates how generating different flow patterns within each incubation unit can accomplish various tasks such as (h) Loading worms, (i) Flushing progeny, and (j) Feeding.

Several aspects of long-term maintenance require daily buffer exchange: worms require fresh food (*E. coli*) and removal of potentially toxic metabolites and old bacteria daily (Figure 1b, c). 200 chambers, each housing an individual worm, each have outlets for loading and daily buffer exchange (Figure 1d). Manual assays require laborious daily transferring of tested populations to new plates to avoid progeny interference with assays (Table 1) or sterilization of adult worms through a variety of methods (fluorodeoxyuridine (FUdR) or blocking of self-sperm development)^22^. However, these progeny-blocking interventions may affect growth, development, aging, or interfere with chemical screening^14,23^. Instead, *Ce*Lab assays of reproductive worms uses flushing of progeny daily. Capable of housing multiple worms (Figure S1a), each incubation arena contains Circular and Progeny chambers that are connected to Perimeter, Progeny, Flush, and Loading outlets (Figure 1b). Sieves at Progeny and Perimeter outlets prevent even the smallest L1 progeny from exiting the incubation arena. However, the sieves at the Flush outlet are big enough for the passage of L1 to L3 animals. Adult worms can only enter through Loading port that is connected to the bottom of a well and can be blocked by a pin (Figure 1c-d). Incubation units are connected in parallel to the chip ports ensuring uniform flow distribution (Figure 1e). Multiple PDMS layers are carefully aligned to create 200 incubation units (Figure S1b-d, Figure S2a-c). On-chip manifolds regulate flow direction through ports of each chip (Figure 1f). Moreover, a valve network controls fluid flow from reservoirs through the main feeding line into inlet chip manifold (Figure 1g). Manipulating these valves can generate flow patterns to carry out various tasks, including loading worms, flushing progeny out of chambers, and feeding or replacing incubation fluid (Figure 1h-j). In *Ce*Lab, setting Perimeter and Progeny ports as inlets, hatched L1-L3 progeny can be cleared from incubation arenas through the flush port in less than 10 minutes with nearly 100% success rate (Figure 1i, Video S1b-c).

#### Re-use of chips

In most microfluidic devices, a new chip must be made for each experiment (Table S1). By contrast, *Ce*Lab chips can be reused upon thorough cleaning. Running dilute chlorine bleach solution for a few minutes can break apart worm and bacterial residue from the chip (Figure 1g) and sterilizes the pipeline, eliminating the risk of contamination for long-term culture.

#### Loading

It is challenging to accurately deliver animals to intended locations in microfluidic chips^24^ (Table S1); the hold-and-burst procedure can take 30 min with 80% yield^25^, or 3 min with 65% yield^26^. Additionally, this highly size-dependent technique is stressful for worms as they are pushed through narrow openings. By contrast, the embedded well-plate feature of the *Ce*Lab chip offers a novel worm-loading technique; loading animals to *Ce*Lab chip is simple and size-independent with 100% yield (Figure 1h, Video S1a). Current worm-loading techniques limit users to test one mutant per chip (Table S1). *Ce*Lab enables the user to load multiple mutants on a chip. After removing pins, wells can be filled in seconds. Deposited animals into wells can be pulled into incubation arenas nearly instantly by applying negative pressure (Figure 1h, Video S1a).

#### Bacterial feeding

Previous designs with continuous bacterial flow forms biofilm streamers that clog chips in a matter of days^27^ (Table S1). Using less sticky *E. coli* strains (HT115) can delay the inevitable^28^, but limits food options. Additionally, significant labor goes into preparing feeding solutions daily. In *Ce*Lab, incubation arenas are filled with feeding solutions once every day, eliminating the risk of clogging (Figure 1j, Video S1d). The bacterial solution is stored in-line and at 4°C for up to two weeks (Figure 1g, 1j). Only 1.5ml of feeding solution is required per chip per day, filling the chip in a matter of seconds (Table 1).

#### Scoring

Some current microfluidic technologies use tracking algorithms for computer-assisted scoring with pre-defined cut-off thresholds, and the quantifications must be reviewed for errors often making them as slow as manual scoring (Table S1). While suitable for scoring motility assays, they are subject to significant limitations, such as successfully identifying bagged or exploded worms. Although *Ce*Lab can be similarly imaged for automated size and motility measurements (Video S1e), it can also be scored manually on a dissecting scope for lifespan, reproductive span, and brood size, closely resembling the operation workflow of plate assays and can be easily adopted by biologists. To expedite manual scoring, we recently developed *Ce*Aid (***C. e****legans* **A**pplication for **i**nputting **d**ata)^29^. Implementing features such as voice command, swiping, and tap gestures, *Ce*Aid does not require the user to shift focus from the chip to the pen and paper throughout the assay (Table 1).

In addition to images, movies of individual wells can be made (Video S2), allowing automated analyses of movement, such as with CeleST^30^, as we have used previously to identify age-related movement disorders^31–33^.

### *Ce*Lab assays recapitulate plate assay results

#### Lifespan assay

Devices that lack pillars often show short lifespans relative to plate assays, indicating that the worms are less healthy in the absence of “dirt” -like conditions^14^. To assess the ability of *Ce*Lab to mimic plate assays, we tested lifespans in the device. *Ce*Lab recapitulates previously-published lifespan results (Figure 2a, c). Moreover, the results are highly reproducible; following step-by-step operation guideline of *Ce*Lab (Figure 1h-j), user-induced variability is eliminated, producing consistent results (Figure 2a, Figure S1e).

**Figure 2:**
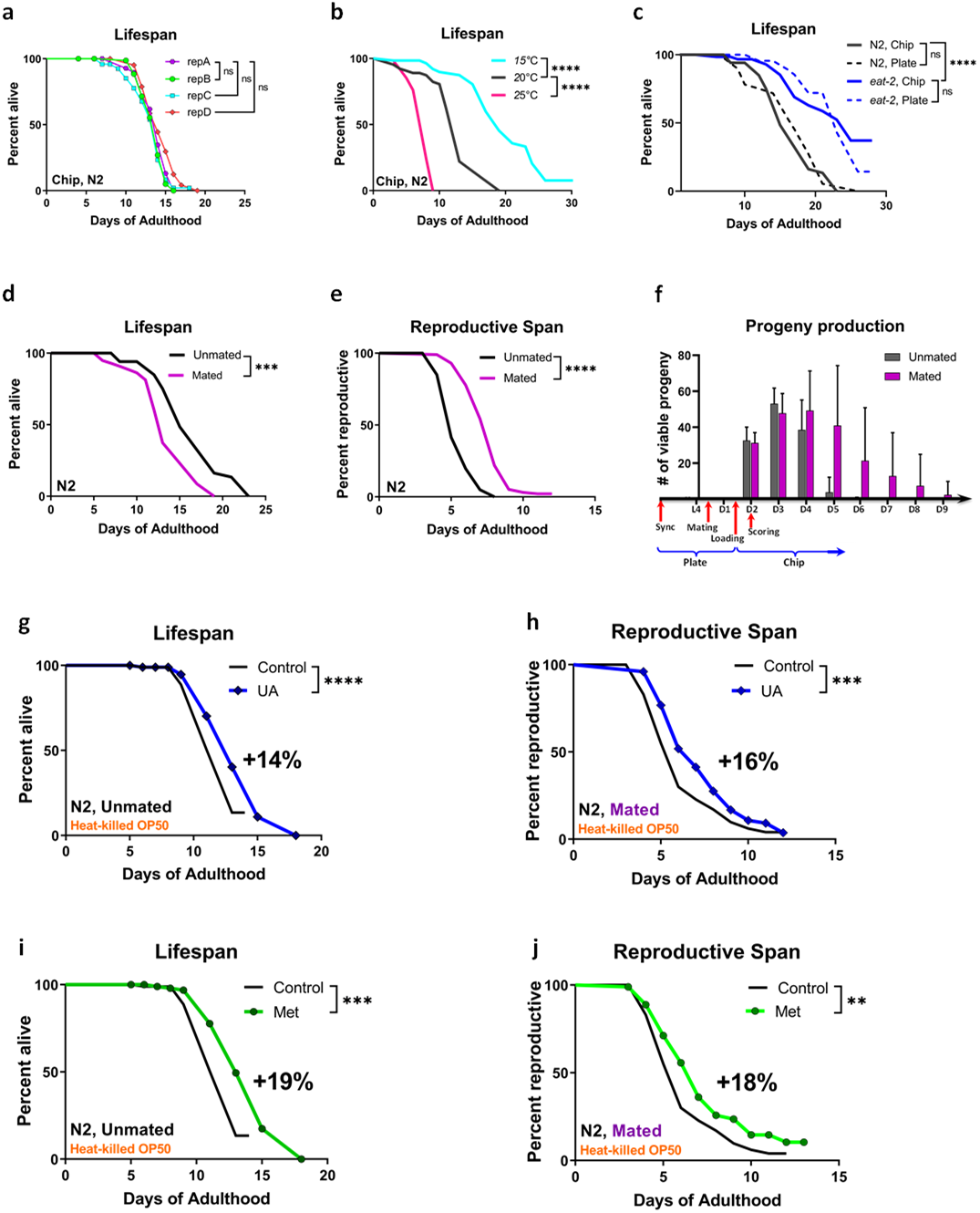
*Ce*Lab reproduces previously-published plate assay results. (a) An example of lifespan data from wild-type (N2) animals; *Ce*Lab’s output data are highly reproducible, (repA, n=99; repB, n=80; repC, n=100; repD, n=66). (b) Wild-type (N2) lifespan at different temperatures. The *Ce*Lab chip replicates the normal lifespan of wild-type N2 worms at different temperatures (15°C, n=66; 20°C, n=66; 25°C, n=66). (c) As shown previously in plate assays, the dietary restriction mutant *eat-2* has an extended lifespan (N2 and *eat-2* on plates, n=80; N2 and *eat-2* in chip, n=66). (d) The *Ce*Lab chip also recapitulates shorter lifespan of mated animals (unmated, n=66; Mated, n=62) (Shi & Murphy, 2014) while (e) reproductive span (Unmated, n=101; Mated, n=100) and (f) progeny production (Unmated, n=51; Mated, n=51) are significantly increased. (g) 50 µM UA treatment in the *Ce*Lab chip replicates previously reported LS extension and (h) reproductive span extension (control, UA, n=100 in both g and h). (i) 50 µM Metformin treatment in the *Ce*Lab chip replicates previously reported LS extension. (j) Mated reproductive span extension is also extended with metformin treatment (control, Metformin, n=100 in both i and j). † indicates loss of hermaphrodites due to matricide. Kaplan-Meier survival tests. NS= not significant. *p < 0.05, **p < 0.01, ***p < 0.001, ****p < 0.0001.

Each *Ce*Lab control center can hold up to 5 chips (Figure 1g, Figure S1c). Chips can be disconnected and incubated at different temperatures. As has been previously shown, worms kept at colder temps live longer (Figure 2b), while worms at higher temperatures live shorter; lifespans performed at these temperatures in *Ce*Lab also show such a temperature difference (Figure 2b).

We also tested the previously-studied dietary restriction longevity mutant *eat-2*, and found that *Ce*Lab recapitulates the extended lifespan previously observed on plates^34^ (Figure 2c). These data also suggest that the worms are not calorically-restricted in the device; in such a case, we would have expected to see a lifespan extension of wild-type worms in the device, and no difference between wild-type worms and *eat-2* mutants in the device.

#### Mated assays

We previously found that while mating extends worms’ reproductive span – because worms are able to use their remaining oocytes once self-sperm are exhausted – mating shortens lifespan^10,35^. Mated worms also produce more progeny later in life^36^. We are able to easily load adult worms and track individuals’ progeny production and health; therefore, worms can be mated prior to loading in the device, and then we simultaneously measure lifespan, reproductive span, and progeny production (Figure 2d-f). It should be noted that not only was *Ce*Lab able to recapitulate the lifespan shortening, reproductive span extension, and progeny production increase upon mating previously shown in plate assays, but *Ce*Lab allows the measurement of these properties in the same individual animals.

#### Reproductive span

We can study reproductive aging by measuring the duration of the mother’s ability to produce offspring daily. To measure reproduction manually, a single fourth larval stage (L4) hermaphrodite is placed on a petri dish with abundant food. Animals are then transferred to a fresh dish daily and are scored for reproductive span or the number of live progeny several days later. Manual reproductive span and brood size assays are highly laborious (Table 1). By contrast, *Ce*Lab is capable of testing 200 worms with any combination of mutants. Thrashing L1 to L3 progeny are easily visible within the incubation arena. During daily inspection, the user scans through the chambers (Video S1e) and records whether individual worms are reproductive by simply swiping up or down on *Ce*Aid. After logging data, hatched progeny are flushed out of the chip. With manual loading and scoring, RS assay using *Ce*Lab is about 7X faster than the manual assay (Table 1). Our *Ce*Lab results recapitulate earlier reproductive span results, including the increase in reproductive span after mating (Figure 2d).

#### Drug treatments

Many drugs and antioxidants have been shown to extend *C. elegans* lifespan^37,38^. Uptake of drugs by animals maintained on plates is an ongoing challenge^39^ and plate assays need to take into account diffusion through the volume of the whole plate. It would be useful to be able to carry out high-throughput drug screens in *C. elegans* in small volumes. Each individual *Ce*Lab chip can hold different experimental fluids (i.e., drugs, RNAi, etc.) with any combination of mutants. *Ce*Lab chips enhance uptake of compounds in pharmacological assays, as animals are submerged in incubation solution. Additionally, only 1.5ml of feeding solution per chip per day is required, reducing the amount of drug required for treatment.

To test the drug screening capability of *Ce*Lab, we measured lifespans and reproductive spans of drug-treated animals. We tracked lifespan of wild-type animals treated with different “longevity” drugs (NMN, NAC, and Urolithin A) (Figure S1g-j; Figure 2g). The *Ce*Lab chip recapitulates the extension of lifespan by NAC and Urolithin A that were previously published^40,41^ (Figure S1g,h; Figure 2g). Additionally, Urolithin A extends mated reproductive span, as we had previously found^42^ (Figure 2h), suggesting that the *Ce*Lab chip will be amenable for drug screening for both lifespan and reproductive spans.

### Metformin treatment increases lifespan and reproductive span

We also tested metformin, the biguanide diabetes drug that has been reported to increase lifespan in a range of organisms, including *C. elegans*^43^; a recent study confirmed that metformin has lifespan and healthspan benefits for several (but not all) *Caenorhabditis* species^44^. A previous study had concluded that metformin’s effects on *C. elegans* lifespan are mediated by perturbation of bacterial folate cycle metabolism, resulting in methionine restriction, rather than through metformin’s effects on the worm itself ^45^. However, those and prior assays were performed with very high (25-100 mM) levels of metformin in the plates. Reasoning that liquid treatment of metformin would be more effective than plate treatment, we reduced the dosage by a thousand-fold from previously published lifespan assays in *C. elegans*^43,45^, and we used the same concentration (50 µM) that we had previously used in a drug screen to reduce Parkinson’s-like phenotypes^31^. Note that in both Mor, et al. (2020) and here, we used heat-killed bacteria to avoid any effects caused by bacterial metabolism. We found that treatment with 50 µM metformin significantly increases both lifespan (+19%, p<0.001) and reproductive span (+18%, p<0.001) (Figure 2i, j). While metformin’s positive effects on lifespan had been reported previously^43^, *Ce*Lab allowed us to observe a simultaneous extension of reproductive span; these effects were observed at 1000-fold lower concentration of metformin than used for previous lifespan assays, but are similar to the positive effects we observed on suppressing Parkinson’s-like behavioral and movement defects at this concentration of metformin^31^.

### *Ce*Lab reduces censored events, thus increasing assay information

One of the problems that plagues *C. elegans* assays, particularly reproductive span assays, is the loss of animals due to matricide (“bagging”), which increases after mating and is exacerbated in many mutant conditions. When an animal bags, it must be censored from lifespan at that point, since it is not considered a natural death. Similarly, the animal should be censored just after bagging in a reproductive span, because the mother is still reproductive at that point but it is impossible to know how much longer she would remain so. Worms are also often censored from assays because they leave the bacterial spot and dry up on the walls of the plate or burrow into the agar (“lost”); some conditions and mutations increase the leaving rate of animals. The combination of leaving and bagging results in high censoring rates in many mated assays performed on plates. However, we find that censoring rates are much lower in the *Ce*Lab chip than on plates (Figure 3a). In addition to eliminating the possibility of leaving and thus reducing censoring due to missing animals, we noticed that worms in the *Ce*Lab chip were far less likely to bag (Figure 3b); this was true both for wild-type worms and for *daf-2* mutants, which generally bag at higher rates than wild-type worms in plate assays (Figure 3b); therefore, we were able to extend the time we can monitor the worms and thus better assess their full reproductive spans and make comparisons between one another. The extended reproductive span of *daf-2* animals is replicated in the *Ce*Lab assay, but with additional time points, as bagging rates are decreased (Figure 3c-d).

**Figure 3.**
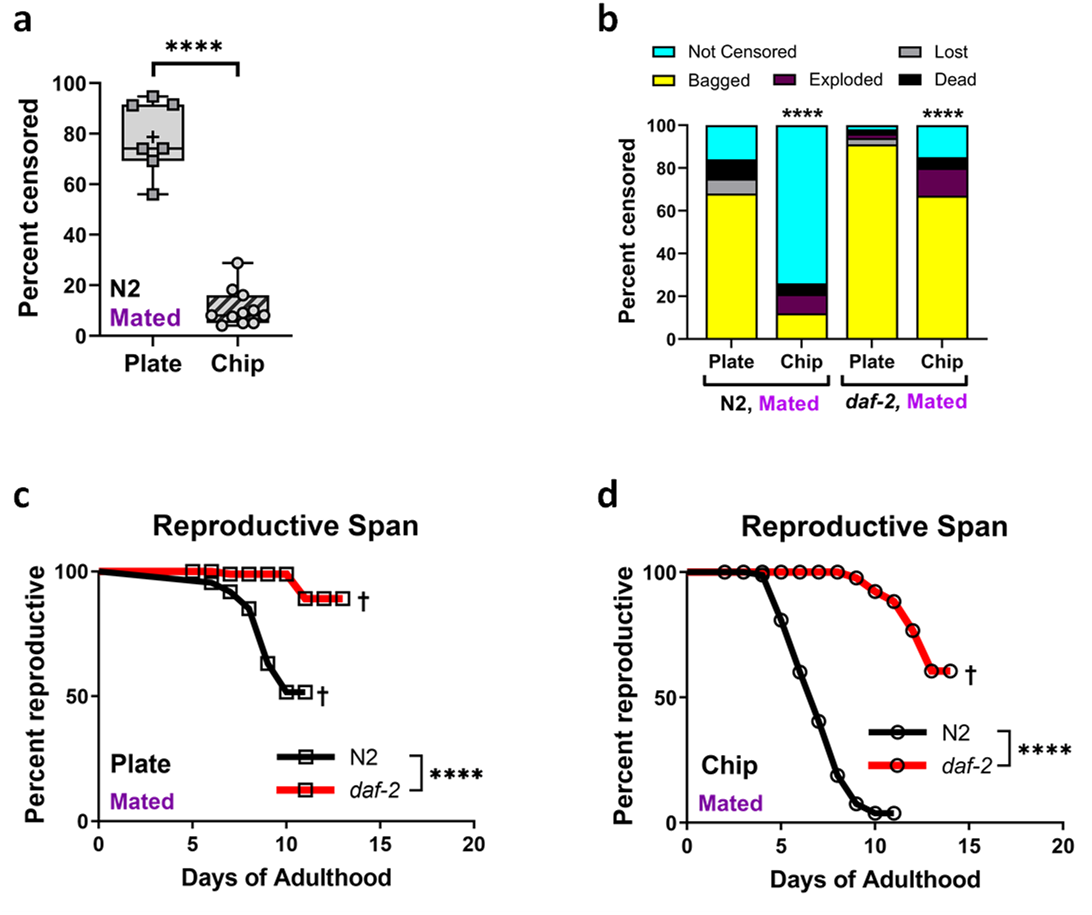
*Ce*Lab reduces censoring rates in reproductive span assays. (a) There are fewer censored events in the *Ce*Lab chip than in plate assays (plate, n=519 from 7 different experiments; Chip, n=852 from 11 different experiments). (b) The *Ce*Lab chip reduces different censoring events, but has the largest effect on the reduction of matricide (bagging, yellow) (Plate: N2, n=89; *daf-2*, n=106; Chip: N2, n=100; *daf-2*, n=196). (c, d) The extended reproductive span of *daf-2* worms is replicated in the *Ce*Lab chip. (Plate: N2, n=89; *daf-2*, n=106; Chip: N2, n=100; *daf-2*, n=196). Kaplan-Meier survival tests. Two-tailed t-tests. Chi-square test. ****p < 0.0001. Box plots show minimum, 25th percentile, median, 75th percentile, maximum.

### Correlational analyses of individual phenotypes

*Ce*Lab can track hundreds of worms, generating both population analyses and data points for each measured individual phenotype (Figure 1g). Simultaneous measurement of visual biomarkers in individuals allows us to uncover new phenotypes and analyze possible correlations between characteristics in individuals, such as size and life history traits that are normally measured in populations (e.g., lifespan and reproductive span). We asked whether a worm’s lifespan is correlated with a worm’s body size, as previous studies have drawn opposite conclusions^46,47^. All body size measurements are done on day 7, which is at the end of the adult growth period^10,46,48^. Different body size parameters are correlated with lifespan in unique ways across the animals tested (Figure 4a, b): while there is no correlation between individual worm length and lifespan of wild type worms, individual *daf-2* worms show a slight positive correlation - that is, longer *daf-2* worms live longer - while longer germlineless *glp-1* animals live shorter (Figure 4a), despite the fact that there is overlap between these two genetic pathways^49^. Carrying out a similar analysis for width, we found that thinner wild-type and *glp-1* animals live longer, while there is no correlation between width and lifespan in *daf-2* (Figure 4b).

**Figure 4:**
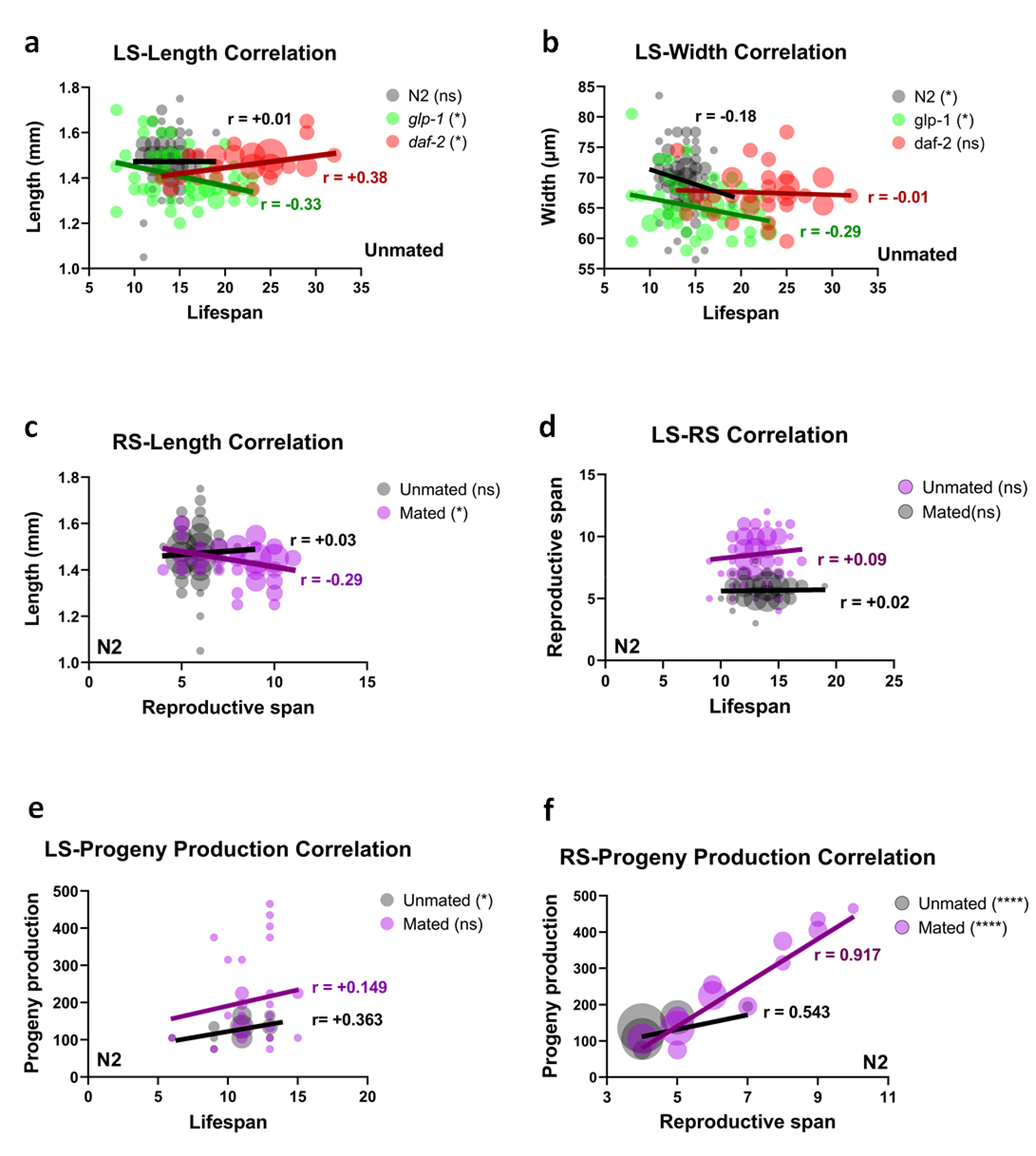
Simultaneous measurement of healthspan biomarkers allows correlation analyses. (a) Length and (b) width correlations with lifespan are different across wild-type animals and longevity and germlineless mutants (N2, n=132 aggregates from 3 different experiments; *glp-1*, n=53 aggregates from 2 different experiments; *daf-2*, n=37). (c) Mated reproductive span is inversely correlated with the worm’s body length (Unmated, n=132 aggregates from 3 different experiments; Mated, n=60 aggregates from 2 different experiments). (d) The length of reproductive span and lifespan is uncoupled for both unmated and mated wild-type animals (Unmated, n=147 aggregates from 3 different experiments; Mated, n=126 aggregates from 3 different experiments). (e) Lifespan is positively correlated with progeny number (Unmated, n=34; Mated, n=34). (f) The number of progeny and the length of reproductive span are strongly correlated, r=0.917, for mated animals (Unmated, n=59; Mated, n=49). Correlation analysis. ns: not significant. *p < 0.05, ****p < 0.0001.

Previously, we had found that mating shortens lifespan and induces shrinking, much of which may be due to water loss^10^; culturing in liquid abrogates water loss-related shrinking (Figure S1f), but lifespan is still shortened post-mating (Figure 2d) likely due to the remaining seminal-fluid-induced pathway^10^. We also found that there is no correlation between length of an individual and the reproductive span in unmated worms. However, mated animals that are shorter in length reproduce longer (Figure 4c).

A prevailing theory of longevity is that there is a trade-off between reproduction and lifespan. According to this theory, high reproduction should have a negative impact on longevity because an organism has limited resources and must choose between investing in germline (progeny production) versus somatic functions (cell repair and thus longevity). This evolutionary trade-off between growth and aging is known as the “Disposable Soma” hypothesis^50^. We find that, for mated and unmated wild-type individuals, lifespan and reproductive span are largely uncoupled (Figure 4d). In other words, there is no relationship between how long an individual animal reproduces with how long or short the animal lives. Moreover, individual lifespan is slightly positively correlated with progeny number (Figure 4e) – that is, the more progeny an animal produces, the longer her lifespan is. Lifespan and reproductive span are uncoupled and further, progeny number may have a positive correlation rather than negative; individuals that produce more progeny actually live slightly longer. Therefore, our data do not support the Disposable Soma hypothesis at the individual level, and instead suggest that an animal that is healthier is more likely to live longer, reproduce longer, and produce more progeny.

Perhaps unsurprisingly, there is a strong correlation between individual mated animals that produce more progeny later in life and length of their reproductive spans (Figure 4f). However, unmated animals also show a positive correlation between reproductive span and progeny number, even though unmated *C. elegans* have a finite number of sperm, suggesting that there may be regulation of the timing of progeny output.

### Heat-killed bacteria extend lifespan and reproductive span

Bacteria can metabolize and thus alter or eliminate chemical compounds, so for many applications, such as drug screens, it is helpful to feed worms a diet of dead bacteria. However, in plate assays, almost all animals leave the heat-killed OP50 *E. coli* bacteria spot and are censored as “lost” (Figure 5a, b), making such screens difficult^51^. By contrast, because worms are retained in their chambers in *Ce*Lab, we can measure their full lifespans on heat-killed bacteria (HK), thus allowing drug screens and other applications that require dead bacteria. We find that unmated worms on an adult-only HK diet (starting at Day 1 of adulthood) have lifespans and reproductive spans that are slightly increased (5%, p<0.001) (Figure 5c, Figure S3a).

**Figure 5:**
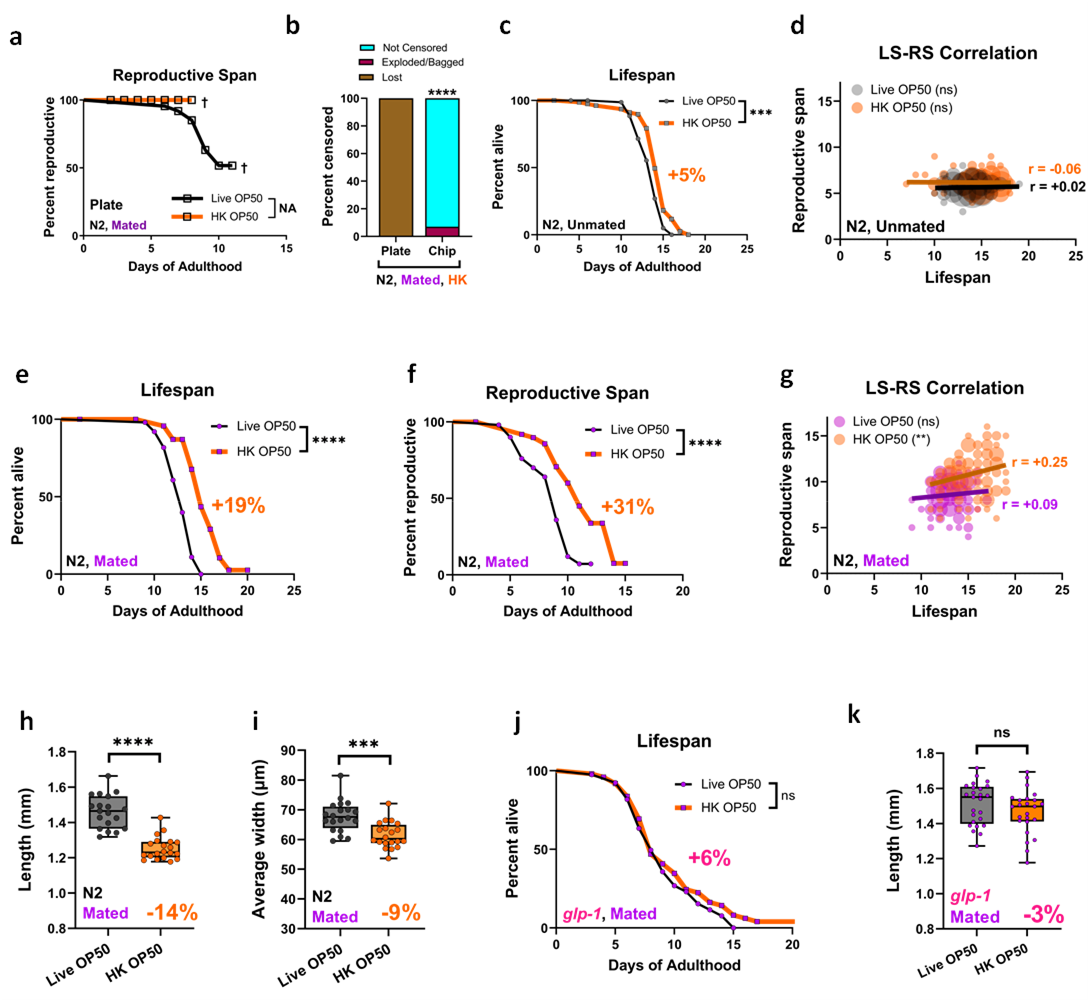
The germline is required for the Heat-killed (HK) bacterial extension of the reproductive span and lifespan. (a) On plate assays, most of the worms are censored due to loss when fed a HK diet (b) but the *Ce*Lab chip reduces this loss (plate, n=100; chip, n=89); (c) Unmated Lifespan is slightly increased on heat-killed bacteria, as measured in the *Ce*Lab chip (Live, n=63; HK, n=50); (d) Lifespan-reproductive span correlation on live and heat-killed bacteria (Live, n=147 aggregates from 3 different experiments; HK, n=168 aggregates from 3 different experiments); (e) Mated lifespan on heat-killed bacteria is significantly increased relative to mated lifespan on live bacteria (Live, n=50; HK, n=50); (f) Mated reproductive span is increased on heat-killed bacteria relative to mated RS on live bacteria (Live, n=50; HK, n=50); (g) On heat-killed bacteria, there is a positive correlation between lifespan and reproductive span (Live, n=126 aggregates from 3 different experiments; HK, n=132 aggregates from 3 different experiments); (h) worms on HK bacteria are significantly shorter (Live, n=19; HK, n=20) and (i) thinner (Live, n=20; HK, n=20); (j) the HK effect requires a functional germline, as *glp-1* mutants do not show a significant increase in lifespan (Live, n=40; HK, n=50); (k) *glp-1* mutants do not shrink on heat-killed bacteria (Live, n=25; HK, n=24). Correlation analysis. Kaplan-Meier survival tests. Two-tailed t-tests. Chi-square test. ns not significant. *p < 0.05, **p < 0.01, ***p < 0.001, ****p < 0.0001. Box plots show minimum, 25th percentile, median, 75th percentile, maximum.

The lifespan difference on the HK diet compared to live bacteria is increased to almost 20% when the worms are mated (Figure 5e); that is, while lifespan is shortened by mating (Shi & Murphy 2014), animals fed heat-killed bacteria suffer less from mating than do worms on live bacteria. Their mated reproductive spans are also extended relative to worms fed live bacteria (Figure 5f). On live bacteria, mating reduces the large difference between lifespan and reproductive span (Figure 2d, e)^10^ so that there is little post-reproductive lifespan. There is also no correlation between lifespan and reproductive span of individuals in unmated conditions and in mated conditions on live bacteria (Figure 5d); however, feeding worms heat-killed bacteria abrogates the lifespan shortening effect of mating. Further, correlation analysis revealed that only mated worms on a HK diet have a positive correlation between lifespan and reproductive span (Figure 5g). That is, mated worms that live longer, reproduce longer if fed a heat-killed bacterial diet. We also found a difference in size between the worms fed a HK diet and a live diet; HK animals are significantly smaller in length and width (Figure 5h-i; Figure S3b-c).

### Worms fed an adult-only heat-killed diet have an extended lifespan and reproductive span that requires the germline

Because we found the most prominent lifespan extension when hermaphrodites are mated with males, and mating increases the amount of progeny produced by 2-3 times^36^ we hypothesized that energy requirements of the germline might activate a stress response or longevity pathway that results in lifespan extension on heat-killed bacteria. That is, we wondered whether the lifespan effect seen in mated worms on heat-killed bacteria is dependent on the germline. The germline proliferation mutant *glp-1* is long-lived^52^, but their lifespan is ∼55% shorter after mating^10^. Since germline proliferation mediates at least part of the lifespan shortening seen in mated worms (Shi & Murphy 2014), it is possible that heat-killed bacteria abrogate the lifespan-shortening effect of mating. To test this possibility, we mated germlineless *glp-1* animals and compared lifespans on live and heat-killed bacteria, and found that lacking a germline abolished the lifespan-extending effect of heat-killed bacteria (Figure 5j). Therefore, germline proliferation is required for the HK diet lifespan extension. Germlineless *glp-1* did not shrink on the HK diet (Figure 5k), suggesting the smaller size induced by HK is mediated by the germline.

### SGK-1 signaling is required for heat-killed bacteria lifespan extension

Since mated hermaphrodites produce significantly more progeny, and mated worms on heat-killed bacteria demonstrate several similarities with dietary restricted (DR) worms (longer lifespan, smaller size; see Figure 5), we hypothesized the DR pathway may be activated by the heat killed bacteria paradigm specifically under mated conditions. DR increases lifespan^34,53–55^ and reproductive span^36,53,56,57^, and is conserved from worms, to flies, to mammals. In worms, the commonly used DR mimic is *eat-2*. We tested *eat-2* mutants in our HK vs. live paradigm; if the DR pathway is activated by HK, then *eat-2* should show no change in lifespan and reproductive span. However, we found that *eat-2* mutants still showed a significant increase in lifespan on heat-killed OP50 (Figure 6a). (Bagging rates, which are already high in *eat-2*, (Figure 6b), limited the ability to complete lifespans.) Therefore, we tested other components of DR. The Nrf2 (NF-E2-related factor)/CNC family of transcription factors ortholog SKN-1 is required for *C. elegans* DR-mediated lifespan increase. Specifically, SKN-1 activity in two sensory neurons is responsible for DR-longevity, and DR-mediated lifespan extension is lost in *skn-1* mutants^58^. Therefore, we tested the transcription factor mutant *skn-1* in our heat-killed vs. live bacteria paradigm. We found that HK increased both lifespan and reproductive span of mated *skn-1* mutants (Figure 6c-d); further, *skn-1* mutants decreased in size on a heat-killed diet (Figure 6e), similar to mated wild-type animals on HK. Therefore, our data suggests HK bacteria-mediated longevity is not dependent on *skn-1*.

**Figure 6:**
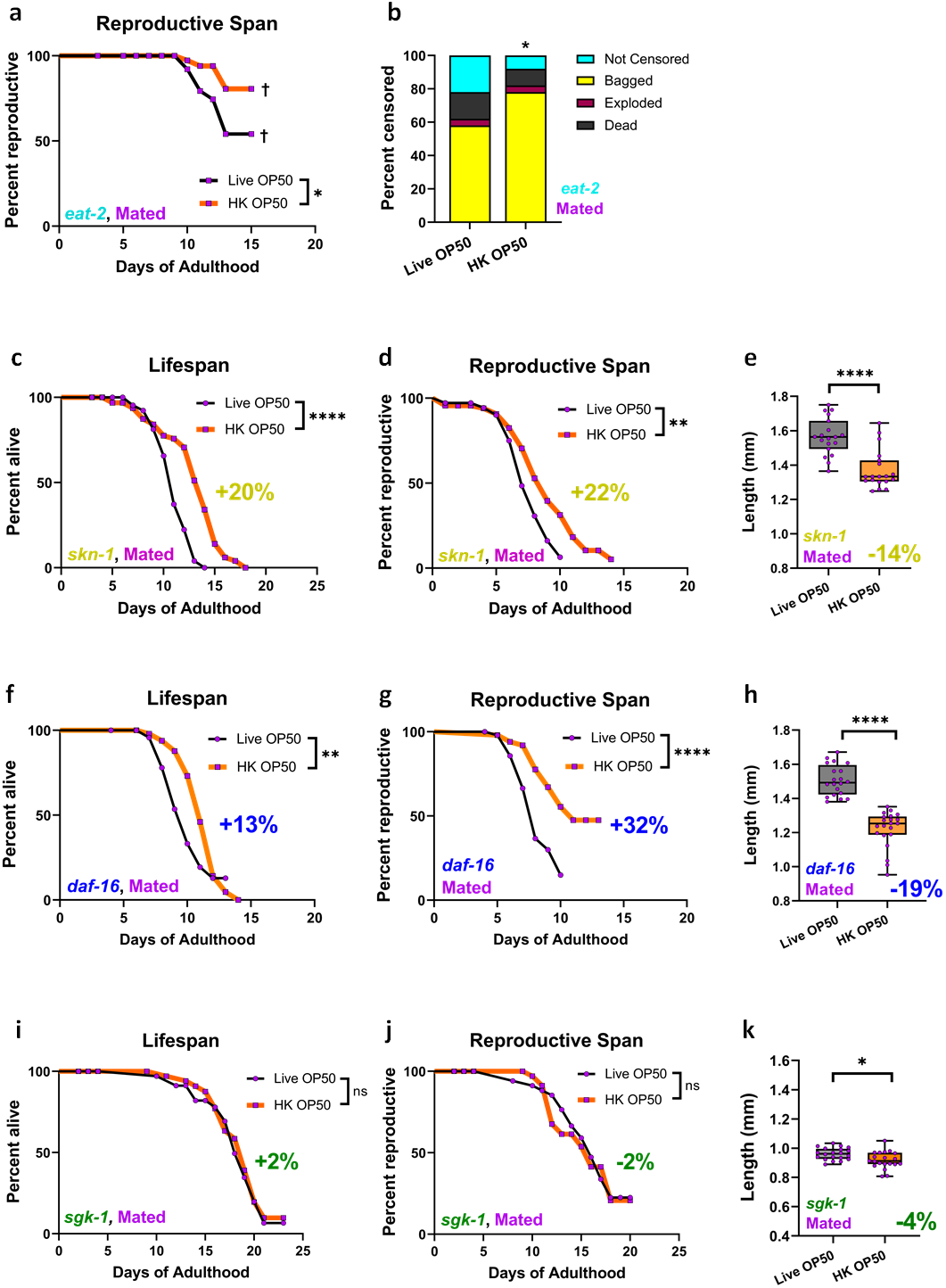
*sgk-1* is required for Heat-killed diet extension. (a) *eat-2* mutants live longer on heat-killed bacteria (Live, n=50) (HK, n=50); (b) bagging rates are high in *eat-2* mutants, and slightly exacerbated on HK; (c) mated *skn-1* mutants still show extended mated lifespan and (d) mated reproductive span on HK (Live, n=59) (HK, n=62); (e) mated *skn-1* mutants shrink on heat-killed bacteria (Live, n=19) (HK, n=19). (f) Mated *daf-16* mutants’ lifespan and (g) reproductive span are increased on HK (Live, n=50) (HK, n=50), and (h) *daf-16* mutants shrink on HK bacteria (Live, n=20) (HK, n=21). (i) Mated *sgk-1* mutants do not live longer or (j) reproduce longer on HK bacteria than on live bacteria (Live, n=40) (HK, n=40). (k) Mated *sgk-1* mutants shrink less on HK (Live, n=20) (HK, n=20). Kaplan-Meier survival tests. Two-tailed t-tests. Chi-square test. ns not significant. *p < 0.05, **p < 0.01, ***p < 0.001, ****p < 0.0001. Box plots show minimum, 25th percentile, median, 75th percentile, maximum.

Next, we tested the Insulin/Insulin-like growth factor 1 (IGF1) pathway. IGF1 signaling mutants have double the lifespan^59^ and reproductive span^56,60^ of wild-type worms through the activation of the FOXO transcription factor DAF-16. If the heat-killed diet activated DAF-16, then *daf-16* might be necessary for the lifespan extension we observe in mated worms on HK. However, mated *daf-16* on HK have both an increased lifespan and reproductive span (Figure 6f, g), and *daf-16* is smaller on heat-killed OP50 (Figure 6h), as we observe in wild-type animals. Together, our data suggests the DAF-2/DAF-16 pathway is not required for the heat-killed lifespan extension paradigm.

We were interested in whether the mTOR pathway might be activated when worms are fed a heat-killed diet. SGK-1 acts downstream of the Rictor/TORC2 complex to mediate nutrient quality signals^61^. *sgk-1* mutants are significantly smaller than wild-type (Figure S3d-e) and have been reported to have shorter lifespans on plates^61,62^ under unmated conditions. However, the lifespan of *sgk-1* mutants in mated conditions has not been previously tested. Unlike mated wild-type, *skn-1*, or *daf-16*, the HK diet was not able to further extend the lifespan of mated *sgk*-*1* animals (Figure 6i). Therefore, we conclude that *sgk-1* is required for the HK-mediated increase in mated lifespan. Additionally, we found that *sgk-1* mutants have an extremely long mated reproductive span on live bacteria (Figure 7b) that not further extended by heat-killed OP50 (Figure 6j); that is, unlike the extension of reproductive span seen in mated wild type, skn-1, and daf-16 animals, HK does not affect sgk-1, suggesting that sgk-1 is required for the HK effect on RS. Finally, mated *sgk-1* mutants shrink less than do mated wild-type, *daf-16*, or *skn-1* worms upon HK feeding (Figure 6k). Taken together, our results suggest that SGK-1 is required for the heat-killed bacterial diet extensions of both lifespan and reproductive span, and that mutation of *sgk-1* essentially phenocopies the effect of HK bacteria on wild-type worms, while reversing the effect of mating on lifespan.

**Figure 7:**
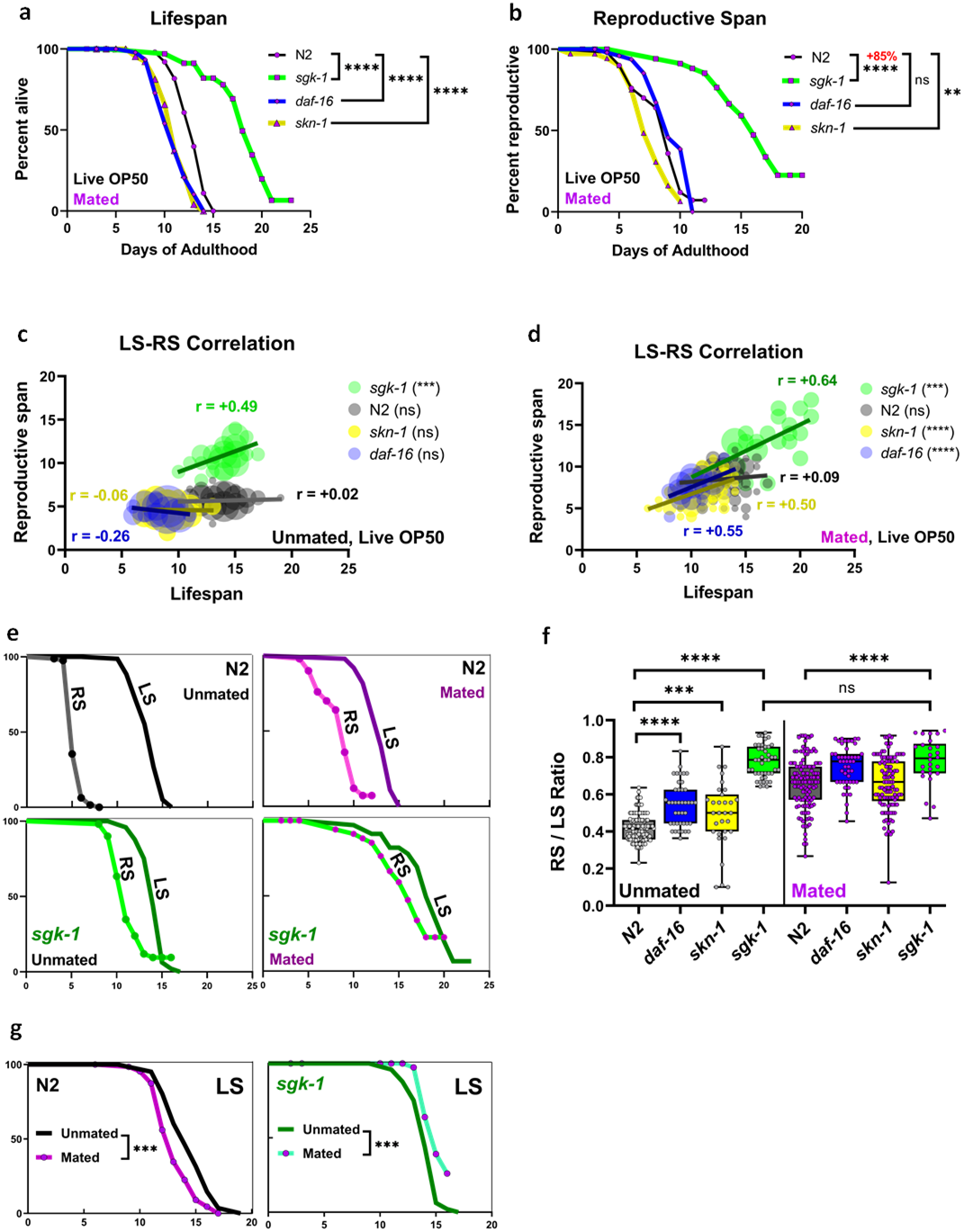
Mated *sgk-1* mutants reproduce nearly until death. Mated *sgk-1* mutants live much longer (a) and reproduce much longer (b) than wild-type (N2), *daf-16*, or *skn-1* mutants, even on live OP50 (N2, n=50) (*sgk-1*, n=40) (*daf-16*, n=50) (*skn-1*, n=70). *sgk-1* mutant individuals with longer reproductive spans live longer under both unmated (c) (*sgk-1*, n=43) (N2, n=147) (*skn-1*, n=28)(*daf-16*, n=46) and mated conditions (d) (*sgk-1*, n=26 aggregates from 2 different experiments) (N2, n=126 aggregates from 3 different experiments) (*skn-1*, n=104 aggregates from 3 different experiments)(*daf-16*, n=48 aggregates from 2 different experiments). (e) Mating reduces the post-reproductive lifespan of wild-type worms, but *sgk-1* mutants extend both lifespan and reproductive span. (f) While the ratio between reproductive span and lifespan is around 0.4 to 0.5 for unmated animals, and rises with mating, *sgk-1* mutants’ RS/LS ratio is not further increased upon mating. (N2-unmated, n=147) (*daf-16*-unmated, n=41) (*skn-1*-unmated, n=28) (*sgk-1*-unmated, n=43) (N2-Mated, n=126) (*daf-16*-Mated, n=48) (*skn-1*-Mated, n=104) (*sgk-1*-Mated, n=26). Kaplan-Meier survival tests. Correlation analysis. One-way ANOVA with Dunnett’s post-hoc. ns not significant. *p < 0.05, **p < 0.01, ***p < 0.001, ****p < 0.0001. Box plots show minimum, 25th percentile, median, 75th percentile, maximum.

### *sgk-1* increases reproductive span of mated animals

To better understand how *sgk-1* mutants affect lifespan and reproductive span after mating on normal live bacteria, we compared them to other well-studied mutants. *daf-16* and *skn-1* mutants have shorter lifespans than mated wild-type worms, but *sgk-1* mated lifespan is much longer (a 40% increase) relative to wild type’s lifespan (Figure 7a). *sgk-1* extends reproductive span even more substantially: mated *sgk-1* animals reproduced 85% longer than wild type (Figure 7b). While the lifespans and reproductive spans of most unmated individual animals are uncorrelated, *sgk-1* mutant individuals’ lifespans and reproductive spans are highly correlated (Figure 7c, d) – that is, an *sgk-1* mutant individual that has an extended lifespan also reproduces for most of its life, even under mated conditions. Most genotypes that are unmated reproduce 40-50% of their lifespan (Figure 7e), but unmated *sgk-1* reproduces nearly until death, thus has a smaller post-reproductive lifespan (Figure 7e, f). In mated animals, all genotypes reproduce until death, including *sgk-1*. Interestingly, there is no difference in the RS/LS ratio between unmated and mated *sgk-1* animals, suggesting SGK-1 plays a role in regulating the length of the reproductive span - essentially “undoing” the deleterious effect of mating (Shi & Murphy, 2014). Wild-type animals have a large post-reproductive lifespan if unmated, and mating simultaneously increases reproductive span and decreases lifespan^10^, reducing the post-reproductive lifespan (Figure 7e-g). Both mated and unmated *sgk-1*, by contrast, reproduce for almost their entire lives (Figure 7e-g). Most surprisingly, while wild-type worms’ mated lifespans are shorter than unmated (Figure 7g; Shi & Murphy), here we found that *skg-1* mutants’ lifespans are almost the same regardless of mating, and even slightly extended upon mating (Figure 7g). Taken together, our data suggest a new role for SGK-1 in the regulation of reproductive span and post-mating lifespan.

## Discussion

Here we have described the design, use, and application of the *Ce*Lab chip for *C. elegans* life history trait measurements, as well as some biological insights gained from experiments done in the chip. We have tried to combine the abilities of other devices to carry out high-throughput monitoring, long-term microfluidic worm incubation, individual worm tracking, and semi-automated measurements with progeny flushing, food replenishment, and some manual options to allow the user flexibility in the types of assays that can be performed. The current design of *Ce*Lab allows us to monitor lifespan, reproductive span, progeny production, and size of hundreds of individuals simultaneously, which enables us to determine correlations that are nearly impossible in lower-throughput, manual population assays. Moreover, the *Ce*Lab chip decreases rates of censoring through bagged and missing worms, enabling us to continue assays much longer than they can be carried out on plates, allowing more complete lifespan and reproductive span analyses.

The ability to examine many individuals for multiple traits simultaneously has allowed us to discover new biological phenomena that open up new research directions. For example, we found a surprising positive correlation between the length of an individual *daf-2* worm and its lifespan. Normally, insulin/IGF-1 signaling reduction is associated with small size and long lifespan, but our data suggest that among isogenic individuals, longer *daf-2* worms will live longer. Yet wild-type worms do not show the same trend; in fact, shorter wild-type individuals live longer; this paradox will require further analyses. Finally, animals that live longer reproduce longer, suggesting that the Disposable Soma theory does not hold true for individual animals. In fact, these findings are reminiscent of the discovery that women who can have children later in life are more likely to live longer^63,64^.

Heat-killed bacteria is a useful food source to avoid bacterial metabolism of drugs when carrying out compound screens, but long-term assays can be difficult due to high rates of censoring due to lawn leaving on plates; *Ce*Lab assay chambers remove this problem. Because we use heat-killed rather than live bacteria for our drug screens, possible side effects from bacterial metabolism are removed. We found that metformin extends lifespan and reproductive span, contrary to a previous report^45^. We had previously found that the same conditions (50 µM metformin on heat-killed bacteria) also suppress a Parkinson’s-like motility phenotype caused by reduction of branched-chain amino acid metabolism^31^. Thus, previously reported effects of metformin on lifespan from live bacteria assays might have been due at least in part to the extremely high metformin concentration used in those studied; in fact, the lifespan shortening observed on 50 mM metformin – 1000x higher than our conditions – is also observed on UV-irradiated *E. coli*^45^, suggesting that this lifespan decrease is caused by the toxic effects of metformin at high concentrations. By contrast, we found that low dosage (50 µM) metformin is beneficial, extending lifespan and reproductive span, in addition to reducing Parkinson’s-like motility defects caused by mitochondrial hyperactivation^31^.

In our studies using heat-killed bacteria, we discovered that both lifespan and reproductive spans of mated animals are extended on an HK diet through a germline- and *sgk-1*-dependent process. Remarkably, *sgk-1* mutants exhibit a phenotype we have not previously observed: not only are their reproductive spans greatly extended after mating relative to wild-type animals, but their lifespan after mating is also greatly extended, reversing the normal lifespan shortening effect of mating (Shi & Murphy, 2014). In fact, *sgk-1* mutants reproduce for almost their entire lifespan, and individuals who live long also reproduce later. These observations would not have been possible in standard plate assays.

Our hope is that the development and adoption of *Ce*Lab will not only increase high-throughput analyses of *C. elegans* life history traits, but also reveal additional new biological findings.

## Materials and Methods

### *C. elegans* strains and maintenance

*C. elegans* strains were grown at 20 °C on nematode growth medium (NGM) plates seeded with OP50 or OP50-1 *Escherichia coli*. The following strains were used in this study: wild-type worms of the N2 Bristol strain, CB1370: daf-2(e1370), CB4037: *glp-1(e2141)*, CF1038: *daf-16(mu86)*, QV225: *skn-1(zj15*), DA465: *eat-2(ad465)*, VC345: *sgk-1(ok538*), CB4108: *fog-2(q71)*. To induce the mutant phenotype, *glp-1(e2141)* was cultured at 25°C until the Day 1 adulthood and shifted back to 20°C to perform the assays. All experiments were performed at 20°C except the ones stated.

### Fabrication

Molds for layer #3 and layer #5 (Figure S1d) are fabricated using SU-8 2075 (MicroChem, Newton, MA, USA) spin-coated at 2150rpm for 30s to create ∼100μm tall patterns (Figure S2a, c). Mold for layer #4 is fabricated using SU-8 2025 (MicroChem, Newton, MA, USA) spin-coated at 4000rpm for 30s to create ∼20μm tall channels (Figure S2b). PDMS (Sylgard 184, Dow Corning Corporation) mixed at 10:1 ratio for all layers, poured on these SU-8 molds, and half-cured at 65°C for 35min. The thickness of layer #1, layer #2, layer#3, layer #4, and layer #5 are 3mm, 0.5mm, 6mm, 3mm, 4mm thickness, respectively (Figure S1d). 600 holes were punched in layer #4 using 0.3mm biopsy punch to create vertical connections for progeny and perimeter outlets (Figure S2b). 200 slanted holes were punched in layer #3 using 0.5mm biopsy punch for worm-loading ports (Figure S2a). 200 holes were punched in thin layer #2 using 1.5mm biopsy punch to create bottom of the wells (Figure 1d). In layer #1, wells are punched using 3mm biopsy punch (Figure 1d). Additionally, 400 holes were punch in layer #1 using 0.75mm biopsy punch to create pin-holders and well to pin-holder connections (Figure S1d). The connection between pin holders and wells are essential as it ensures sterilization of pin holders when wells are filled with bleach. Finally, progeny, perimeter, and flush ports in layer #3 and layer #4 are punched with 1mm biopsy punch. Half-cured layers are stacked and aligned with high precision with 15µm tolerance. Stacked layers are then cured at 65°C for 24hr.

### Synchronizing

Gravid hermaphrodites are treated with a 15% hypochlorite solution (e.g., 8.0 mL water, 0.5 mL 5N KOH, 1.5 mL sodium hypochlorite), followed by multiple rounds of washing collected eggs in M9 buffer.

### Mating

Animals at the L4 stage were mated with *fog-2(q71)* males at a 3:1 ratio for 24 hrs from L4 to Day 1 of adulthood, then were singled into the wells of *Ce*Lab chips on Day 1 of adulthood.

### Loading animals

Spraying top wells with ethanol the day prior to loading worms is also essential to make sure no contamination will be introduced to the chip. A dilute solution of feeding solution (6 OD600) should be used during loading to prevent starvation. Pins are then moved from worm-loading ports to pin-holders. Positive pressure can fill all wells with diluted OP50 solution in a matter of seconds (Figure 2b). Animals are then deposited into wells. By applying negative pressure, all worms are pulled into incubation chambers instantly (Video S1a).

### Preparing the feeding solution

Animals and their progeny should have enough food between feeding sessions. It is previously shown that 10 adult worms consume ∼1.5×10^7^ colony-forming units (cfu)/ml over 72hr (D1:D4)^65^. We calculated that the required concentration of bacterial solution for incubating one D1 adult worm overnight per chamber is 60 OD600 (1 OD600 corresponds to 1.5×10^8^ cfu/ml). The feeding solution is prepared by spinning down cultivated LB broth at 4500 RPM for 30min and resuspending the pellet in S-complete medium. The feeding solution must then be filtered through a 10µm cell strainer. The OD600 measurement using spectrophotometer should be carried out with a 1:100 diluted sample so that the measured optical density is within 0-1. OD600 was measured by a Nanodrop 2000C (Thermo Scientific). The prepared bacterial solution can then be gradually diluted with S-medium as progeny production decrease during the experiment. When animals stop reproducing, chips can be flushed every 2-3 days and then loading with 10 OD600 feeding solutions.

### Heat-killed OP50

OP50 bacteria was killed by incubation at 65 °C for 30 min.

### Scoring using *Ce*Aid

To expedite manual scoring, we recently developed *Ce*Aid (***C. e****legans* **A**pplication for **i**nputting **d**ata) ^29^. Implementing features such as voice command, swiping, and tap gestures, *Ce*Aid does not require the user to shift focus from the chip to the pen and paper throughout the assay.

### Reproductive span assay

Thrashing L1 to L3 progeny are easily visible within the incubation arena. Reproductive cessation was defined as the last day of progeny production^66^. Worms were censored on the day of matricide (bagging is defined as progeny hatching within mother), abnormal vulva structures, and dead.

### Lifespan assay

Animals were scored daily and defined as dead when they no longer move or responded to touch^59^. Similar to gently touching worms with the pick in plate LS assays, user can flick pins to stimulate worms. worms were censored on the day of matricide (bagging is defined as progeny hatching within mother), abnormal vulva structures.

### Progeny production

Every 24 hours before flushing, a 3s video of each incubation arenas were captured. These videos were then analyzed to score for numbers of hatched progeny.

### Flushing

After logging data, flushing protocol can be initiated to remove progeny and/or provide a fresh food source (Figure 1i and Video S1b-c). Moreover, in cases of bagged worms where progeny grow beyond L3 within the mother, pins can be removed from worm-loading port and L4 progeny is flushed into the well. In rare cases where the flow to a channel is blocked, a gentle push on progeny chamber can simply restore the flow without hurting adult worms as hexagonally arranged pillars are highly effective in protecting animals within the circular chamber.

### Feeding

*Ce*Lab, incubation arenas are filled with feeding solutions once every day (Figure 1j and Video S1d). Bacterial solution is stored in-line and at 4°C for up to two weeks (Figure 1g and Figure S1c). Only 1.5ml of feeding solution is required per chip per day filling chip in matter of seconds. All valves of on-chip manifolds are then closed for overnight incubation.

### Cleaning chips

Running dilute chlorine bleach solution for a few minutes can break apart worm and bacterial residues from the chip. It also sterilizes the pipeline, eliminating the risk of contamination for long-term culture. After exposure to bleach, the chip should be rinsed with DI water for 15 minutes.

### Drug treatment

NAC, NMN, Urolithin A, and Metformin were obtained through Fisher Scientific and diluted in DMSO. The control is composed of the incubation solution with 0.3% v/v DMSO, and the experimental incubation solution has 0.3% v/v DMSO and 50 µM Urolithin A, 5 mM NAC, 5 mM NMN or 50 μM Metformin. All drug treatments were started on day 1 of adulthood.

### Body size measurement

Images were taken on day 7 with an ocular-fitted phone camera attached to a standard dissection microscope (Video S1e). For body area measurement, the polygon selection tool was used to outline the boundary of each worm. For body length measurement, the middle line of each worm was delineated using the segmented line tool. Average width is calculated by dividing body area by length.

#### Statistics and reproducibility

Reproductive span and lifespans assays were assessed using the standard Kaplan-Meier log rank survival tests (Table S2A). Censoring rate analyses used chi square test to determine whether there was any significant difference between populations for each category (Table S2B). Correlation analysis is used to statistically analyze the relationship between different healthspan biomarkers (Table S2C). For comparisons between two groups, an unpaired Student’s t test was performed. For comparisons between multiple groups, One-Way ANOVA was performed with post-hoc testing. Replicates for experiments were carried out on separate days with separate, independent populations. GraphPad Prism was used for all statistical analyses.

## Acknowledgments

We thank *the C. elegans* Genetics Center for strains and the Murphy lab for discussion. This work was supported by a GCRLE award to CTM (GCRLE-0220), DP1 Pioneer NIH award to CTM (1DP1AG077430-01), TR01 award to CTM (5R01AT011963-02), and Glenn Foundation for Medical Research award. CTM is the Director of the Glenn Center for Aging Research at Princeton and the Director of the Simons Collaboration on Plasticity in the Aging Brain.

## References

1. Saberi-Bosari, S., Huayta, J. & San-Miguel, A. A microfluidic platform for lifelong high-resolution and high throughput imaging of subtle aging phenotypes in C. elegans. Lab on a Chip 18, 3090–3100 (2018).

2. Churgin, M. A. et al. Longitudinal imaging of Caenorhabditis elegans in a microfabricated device reveals variation in behavioral decline during aging. eLife 6, e26652 (2017).

3. Stroustrup, N. et al. The C. elegans Lifespan Machine. Nat Methods 10, 665–670 (2013).

4. Coleman-Hulbert, A. L. et al. Caenorhabditis Intervention Testing Program: the creatine analog β-guanidinopropionic acid does not extend lifespan in nematodes. microPublication Biology (2020).

5. Lucanic, M. et al. Impact of genetic background and experimental reproducibility on identifying chemical compounds with robust longevity effects. Nat Commun 8, 14256 (2017).

6. Morshead, M. L. et al. Caenorhabditis Intervention Testing Program: the farnesoid X receptor agonist obeticholic acid does not robustly extend lifespan in nematodes. MicroPubl Biol 2020, 10.17912/micropub.biology.000257 (2020).

7. Kerr, R. A., Roux, A. E., Goudeau, J. & Kenyon, C. The C. elegans Observatory: High-throughput exploration of behavioral aging. Front Aging 3, 932656 (2022).

8. Hahm, J.-H. et al. C. elegans maximum velocity correlates with healthspan and is maintained in worms with an insulin receptor mutation. Nat Commun 6, 8919 (2015).

9. Li, S., Stone, H. A. & Murphy, C. T. A microfluidic device and automatic counting system for the study of C. elegans reproductive aging. Lab Chip 15, 524–531 (2015).

10. Shi, C. & Murphy, C. T. Mating Induces Shrinking and Death in Caenorhabditis Mothers. Science 343, 536–540 (2014).

11. Ghosh, R. & Emmons, S. W. Episodic swimming behavior in the nematode C. elegans. Journal of Experimental Biology 211, 3703– 3711 (2008).

12. Ikeda, Y. et al. C. elegans episodic swimming is driven by multifractal kinetics. Scientific reports 10, 1–17 (2020).

13. Ben-Yakar, A. High-content and high-throughput in vivo drug screening platforms using microfluidics. Assay and drug development technologies 17, 8–13 (2019).

14. Rahman, M. et al. NemaLife chip: a micropillar-based microfluidic culture device optimized for aging studies in crawling C. elegans. Scientific reports 10, 1–19 (2020).

15. Atakan, H. B. et al. Automated platform for long-term culture and high-content phenotyping of single C. elegans worms. Scientific reports 9, 1–14 (2019).

16. Cornaglia, M. et al. An automated microfluidic platform for C. elegans embryo arraying, phenotyping, and long-term live imaging. Scientific reports 5, 1–13 (2015).

17. Gokce, S. K. et al. A fully automated microfluidic femtosecond laser axotomy platform for nerve regeneration studies in C. elegans. PloS one 9, e113917 (2014).

18. Albrecht, D. R. & Bargmann, C. I. High-content behavioral analysis of Caenorhabditis elegans in precise spatiotemporal chemical environments. Nature methods 8, 599–605 (2011).

19. Lee, H. et al. A multi-channel device for high-density target-selective stimulation and long-term monitoring of cells and subcellular features in C. elegans. Lab on a Chip 14, 4513–4522 (2014).

20. Mondal, S. et al. Large-scale microfluidics providing high-resolution and high-throughput screening of Caenorhabditis elegans poly-glutamine aggregation model. Nature communications 7, 1–11 (2016).

21. Lockery, S. R. et al. Artificial dirt: microfluidic substrates for nematode neurobiology and behavior. Journal of neurophysiology 99, 3136–3143 (2008).

22. Midkiff, D. & San-Miguel, A. Microfluidic technologies for high throughput screening through sorting and on-chip culture of C. elegans. Molecules 24, 4292 (2019).

23. Angeli, S. et al. A DNA synthesis inhibitor is protective against proteotoxic stressors via modulation of fertility pathways in Caenorhabditis elegans. Aging (Albany NY) 5, 759 (2013).

24. Banse, S. A., Blue, B. W., Robinson, K. J., Jarrett, C. M. & Phillips, P. C. The Stress-Chip: A microfluidic platform for stress analysis in Caenorhabditis elegans. PLoS One 14, e0216283 (2019).

25. Le, K. N. et al. An automated platform to monitor long-term behavior and healthspan in Caenorhabditis elegans under precise environmental control. Communications biology 3, 1–13 (2020).

26. Chung, K. et al. Microfluidic chamber arrays for whole-organism behavior-based chemical screening. Lab on a Chip 11, 3689–3697 (2011).

27. Drescher, K., Shen, Y., Bassler, B. L. & Stone, H. A. Biofilm streamers cause catastrophic disruption of flow with consequences for environmental and medical systems. Proceedings of the National Academy of Sciences 110, 4345–4350 (2013).

28. Letizia, M. C. et al. Microfluidics-enabled phenotyping of a whole population of C. elegans worms over their embryonic and post-embryonic development at single-organism resolution. Microsystems & nanoengineering 4, 1–11 (2018).

29. Sohrabi, S., Moore, R. S. & Murphy, C. T. Ce Aid: a smartphone application for logging and plotting Caenorhabditis elegans assays. G3 11, jkab259 (2021).

30. Restif, C. et al. CeleST: Computer Vision Software for Quantitative Analysis of C. elegans Swim Behavior Reveals Novel Features of Locomotion. PLOS Computational Biology 10, e1003702 (2014).

31. Mor, D. E. et al. Metformin rescues Parkinson’s disease phenotypes caused by hyperactive mitochondria. Proceedings of the National Academy of Sciences 117, 26438–26447 (2020).

32. Sohrabi, S., Mor, D. E., Kaletsky, R., Keyes, W. & Murphy, C. T. High-throughput behavioral screen in C. elegans reveals Parkinson’s disease drug candidates. Commun Biol 4, 1–9 (2021).

33. Yao, V. et al. An integrative tissue-network approach to identify and test human disease genes. Nat Biotechnol 10.1038/nbt.4246 (2018) doi:10.1038/nbt.4246.

34. Lakowski, B. & Hekimi, S. The genetics of caloric restriction in Caenorhabditis elegans. Proceedings of the National Academy of Sciences 95, 13091–13096 (1998).

35. Gems, D. & Riddle, D. L. Genetic, behavioral and environmental determinants of male longevity in Caenorhabditis elegans. Genetics 154, 1597–1610 (2000).

36. Hughes, S. E., Evason, K., Xiong, C. & Kornfeld, K. Genetic and pharmacological factors that influence reproductive aging in nematodes. PLoS Genet preprint, e25 (2005).

37. Petrascheck, M., Ye, X. & Buck, L. B. An antidepressant that extends lifespan in adult Caenorhabditis elegans. Nature 450, 553– 556 (2007).

38. Petrascheck, M., Ye, X. & Buck, L. B. A high-throughput screen for chemicals that increase the lifespan of Caenorhabditis elegans. Annals of the New York Academy of Sciences 1170, 698–701 (2009).

39. Evason, K., Huang, C., Yamben, I., Covey, D. F. & Kornfeld, K. Anticonvulsant medications extend worm life-span. science 307, 258–262 (2005).

40. Oh, S.-I., Park, J.-K. & Park, S.-K. Lifespan extension and increased resistance to environmental stressors by N-Acetyl-L-Cysteine in Caenorhabditis elegans. Clinics (Sao Paulo) 70, 380– 386 (2015).

41. Ryu, D. et al. Urolithin A induces mitophagy and prolongs lifespan in C. elegans and increases muscle function in rodents. Nature Medicine 22, 879–888 (2016).

42. Cota, V., Sohrabi, S., Kaletsky, R. & Murphy, C. T. Oocyte mitophagy is critical for extended reproductive longevity. Plos Genetics 18, e1010400 (2022).

43. Onken, B. & Driscoll, M. Metformin induces a dietary restriction– like state and the oxidative stress response to extend C. elegans healthspan via AMPK, LKB1, and SKN-1. PloS one 5, e8758 (2010).

44. Onken, B. et al. Metformin treatment of diverse Caenorhabditis species reveals the importance of genetic background in longevity and healthspan extension outcomes. Aging Cell 21, e13488 (2022).

45. Cabreiro, F. et al. Metformin retards aging in C. elegans by altering microbial folate and methionine metabolism. Cell 153, 228–239 (2013).

46. Pincus, Z., Smith-Vikos, T. & Slack, F. J. MicroRNA predictors of longevity in Caenorhabditis elegans. PLoS genetics 7, e1002306 (2011).

47. Zhang, W. B. et al. Extended twilight among isogenic C. elegans causes a disproportionate scaling between lifespan and health. Cell Systems 3, 333–345 (2016).

48. Hulme, S. E. et al. Lifespan-on-a-chip: microfluidic chambers for performing lifelong observation of C. elegans. Lab on a Chip 10, 589–597 (2010).

49. Hsin, H. & Kenyon, C. Signals from the reproductive system regulate the lifespan of C. elegans. Nature 399, 362–366 (1999).

50. Kirkwood, T. B. Evolution of ageing. Nature 270, 301–304 (1977).

51. Qi, B., Kniazeva, M. & Han, M. A vitamin-B2-sensing mechanism that regulates gut protease activity to impact animal’s food behavior and growth. Elife 6, e26243 (2017).

52. Arantes-Oliveira, N., Apfeld, J., Dillin, A. & Kenyon, C. Regulation of Life-Span by Germ-Line Stem Cells in Caenorhabditis elegans. Science 295, 502–505 (2002).

53. Chapman, T. & Partridge, L. Female fitness in Drosophila melanogaster: an interaction between the effect of nutrition and of encounter rate with males. Proceedings of the Royal Society of London. Series B: Biological Sciences 263, 755–759 (1996).

54. Klass, M. R. Aging in the nematode Caenorhabditis elegans: Major biological and environmental factors influencing life span. Mechanisms of Ageing and Development 6, 413–429 (1977).

55. McCay, C. M., Crowell, M. F. & Maynard, L. A. The Effect of Retarded Growth Upon the Length of Life Span and Upon the Ultimate Body Size: One Figure. The Journal of Nutrition 10, 63– 79 (1935).

56. Huang, C., Xiong, C. & Kornfeld, K. Measurements of age-related changes of physiological processes that predict lifespan of Caenorhabditis elegans. Proceedings of the National Academy of Sciences 101, 8084–8089 (2004).

57. Selesniemi, K., Lee, H.-J. & Tilly, J. L. Moderate caloric restriction initiated in rodents during adulthood sustains function of the female reproductive axis into advanced chronological age. Aging Cell 7, 622–629 (2008).

58. Bishop, N. A. & Guarente, L. Two neurons mediate diet-restriction-induced longevity in C. elegans. Nature 447, 545–549 (2007).

59. Kenyon, C., Chang, J., Gensch, E., Rudner, A. & Tabtiang, R. A C. elegans mutant that lives twice as long as wild type. Nature 366, 461–464 (1993).

60. Luo, S., Kleemann, G. A., Ashraf, J. M., Shaw, W. M. & Murphy, C. T. TGF-β and Insulin Signaling Regulate Reproductive Aging via Oocyte and Germline Quality Maintenance. Cell 143, 299–312 (2010).

61. Soukas, A. A., Kane, E. A., Carr, C. E., Melo, J. A. & Ruvkun, G. Rictor/TORC2 regulates fat metabolism, feeding, growth, and life span in Caenorhabditis elegans. Genes & development 23, 496–511 (2009).

62. Chen, A. T.-Y., Guo, C., Dumas, K. J., Ashrafi, K. & Hu, P. J. Effects of Caenorhabditis elegans sgk-1 mutations on lifespan, stress resistance, and DAF-16/FoxO regulation. Aging Cell 12, 932– 940 (2013).

63. Perls, T. T., Alpert, L. & Fretts, R. C. Middle-aged mothers live longer. Nature 389, 133–133 (1997).

64. Sun, F. et al. Extended maternal age at birth of last child and women’s longevity in the Long Life Family Study. Menopause 22, 26 (2015).

65. Gomez-Amaro, R. L. et al. Measuring food intake and nutrient absorption in Caenorhabditis elegans. Genetics 200, 443–454 (2015).

66. Luo, S., Shaw, W. M., Ashraf, J. & Murphy, C. T. TGF-ß Sma/Mab signaling mutations uncouple reproductive aging from somatic aging. PLoS genetics 5, e1000789 (2009).

