## Supplementary information for "*Ce*Lab, a Microfluidic Platform for the Study of Life History Traits, reveals Metformin and SGK-1 regulation of Longevity and Reproductive Span"

**Table S1: Leading technologies developed to automate *C. elegan*s assays.** All use microfluidic technology except the shaded rows, some of which are novel incubation devices and others are modified plate assays. Lifespan (LS), Dietary Restriction (DR), Reproductive span (RS).

| Reference | Assay/Ability | Worms per Unit | # of Units | Advantages | Limitations |
| --- | --- | --- | --- | --- | --- |
| Le, et al. (2020)  HeALTH | LS (DR) | 1 | 60 | - Multiple chips are monitored simultaneously on linear stage | - No pillars in chambers  - Complicated to load worms  - Difficult to operate |
| Rahman, et al. (2020)  NemaLife | LS (DR); Motility; Food intake | 15 | 9 | - Intermittent fluid flow  - Tests multiple populations | - No individual tracking  - Limited capacity |
| Banse, et al. (2019)  Stress-Chip | LS (DR); Oxidative and osmotic stress | 1 | 100 | - Automated survival scoring | - Complicated to load worms  - Prone to clogging  - One population per chip |
| Letizia, et al. (2018) | Developmental rate | 1 | 16 | - Great tool for embryonic and post-embryonic studies | - Application specific  - Limited capacity  - One population per chip |
| Mondal, et al. (2016) | High-throughput imaging | 40 | 96 | - High-resolution and high-speed image acquisition  - Tests multiple populations | - Application specific |
| Li, et al. (2015) | RS; Brood size | 1 | 16 | - Automated RS and brood size scoring | - Application specific  - Complicated to load animals  - Small number of worms |
| Xian, et al. (2013)  WormFarm | LS (DR) | 40 | 8 | - Automated survival, body size and motility scoring  - Tests multiple populations | - No pillars in chambers  - Difficult to operate  - No individual tracking |
| Albrecht, et al. (2011) | Chemosensory behaviors | 25 | 1 | - Great tool for chemosensory stimulus studies | - Application specific |
| Churgin, et al. (2017)  WormMotel | LS; Motility | 1 | 240 | - Automated survival and motility analysis of individual worms | - No progeny removal mechanism  - Laborious to fabricate |
| Pincus, et al. (2011) | LS; Motility;  High-throughput imaging | 1 | 100 | - Enables long-term tracking of individual worms | - No progeny removal mechanism  - No mid-life manipulation |
| Stroustrup, et al. (2013)  Lifespan Machine | LS | 35 | 10×16 | - Automated data acquisition and processing | - No progeny removal mechanism  - No individual tracking |
| Kerr, et al. (2022)  *C. elegans* Observatory | LS; Body size; Motility | 40-60 | 64×9 | - Automated data acquisition and processing | - No progeny removal mechanism  - No individual tracking |

**Table S2: Statistical data tables****.** (A) Lifespan, reproductive span, progeny production, length, width, and LS/RS ratio statistical information. (B) Censoring rate statistical information. (C) Correlation analysis statistical information. Statistics were performed on Prism.

**Table A.**

| **Fig.** | **Assay** | **Test Condition** | **Comparison** | **Rep #** | **Mean** | **% Change** | **p-value** | **n** |
| --- | --- | --- | --- | --- | --- | --- | --- | --- |
| 2a | LS | N2, Unmated, Live OP50, Chip | repB/repA | NA | 13.46/13.63 | -1.24 | 0.2074, n.s. | 80/99 |
|  |  |  | repC/repA | NA | 13.04/13.63 | -4.32 | 0.1187, n.s. | 100/99 |
|  |  |  | repD/repA | NA | 14.27/13.63 | +4.48 | 0.0587, n.s. | 66/99 |
| 2b | LS | N2, Unmated, Live OP50, Chip | 15°C/20°C | 1 | 19.81/12.19 | +62.51 | <0.0001 | 66/66 |
|  |  |  | 25°C/20°C | 1 | 7.25/12.19 | -40.52 | <0.0001 | 66/66 |
| 2c | LS | N2, Unmated, Live OP50 | Chip/Plate | 1 | 16.28/16.70 | -2.51 | 0.253, n.s. | 66/80 |
|  |  | *eat-2,* Unmated, Live OP50 |  | 1 | 22.24/22.97 | -3.17 | 0.135, n.s. | 66/80 |
|  |  | Unmated, Live OP50, Chip | *eat-2*/N2 | 1 | 22.24/16.28 | +36.60 | <0.0001 | 66/66 |
| 2d | LS | N2, Live OP50, Chip | Mated/Unmated | 1 | 13.33/16.28 | -18.12 | 0.0003 | 62/66 |
|  |  |  |  | 2 | 13.08/14.35 | -8.85 | 0.0006 | 67/66 |
|  |  |  |  | 3 | 11.77/13.04 | -9.22 | <0.0001 | 100/100 |
| 2e | RS | N2, Live OP50, Chip | Mated/Unmated | 1 | 7.56/5.51 | +37.20 | <0.0001 | 100/101 |
|  |  |  |  | 2 | 9.51/5.87 | +62.01 | <0.0001 | 67/66 |
|  |  |  |  | 3 | 7.93/5.84 | +35.78 | <0.0001 | 98/99 |
| 2f | PP | N2, Chip | Mated/Unmated | 1 | NA | NA | NA | 51/51 |
| 2g | LS | N2, Unmated, HK OP50 | UA/Control | 1 | 13.40/11.74 | +14.14 | 0.0001 | 100/100 |
| 2h | RS | N2, Mated, HK OP50 | UA/Control | 1 | 7.30/6.27 | +16.34 | 0.0036 | 100/100 |
|  |  |  |  | 2 | 7.68/6.97 | +10.18 | 0.0006 | 100/100 |
| 2i | LS | N2, Unmated, HK OP50 | Met/Control | 1 | 13.97/11.74 | +19.00 | <0.0001 | 100/100 |
|  |  |  |  | 2 | 14.30/12.82 | +11.54 | <0.0001 | 100/100 |
| 2j | RS | N2, Mated, HK OP50 | Met/Control | 1 | 7.40/6.27 | +17.90 | 0.0026 | 100/100 |
|  |  |  |  | 2 | 12.22/10.67 | +14.52 | 0.0002 | 99/95 |
| 3c | RS | Mated, Live OP50, Plate | *daf-2*/N2 | 1 | 12.74/9.87 | +29.07 | <0.0001 | 106/89 |
| 3d | RS | Mated, Live OP50, Chip | *daf-2*/N2 | 1 | 13.15/7.10 | +85.20 | <0.0001 | 196/100 |
| 5a | RS | N2, Mated, Plate | HK/Live OP50 | 1 | NA/9.87 | NA | NA | 100/89 |
| 5c | LS | N2, Unmated, Chip | HK/Live OP50 | 1 | 14.20/13.46 | +5.49 | 0.0003 | 80/80 |
|  |  |  |  | 2 | 14.18/14.35 | -1.18 | 0.7757, n.s. | 50/66 |
|  |  |  |  | 3 | 15.99/14.52 | +10.12 | <0.0001 | 50/40 |
| 5e | LS | N2, Mated, Chip | HK/Live OP50 | 1 | 15.25/12.83 | +18.86 | <0.0001 | 50/50 |
|  |  |  |  | 2 | 16.54/13.08 | +26.45 | <0.0001 | 50/67 |
|  |  |  |  | 3 | 15.87/13.41 | +18.34 | <0.0001 | 50/40 |
| 5f | RS | N2, Mated, Chip | HK/Live OP50 | 1 | 11.18/8.53 | +31.06 | <0.0001 | 50/50 |
|  |  |  |  | 2 | 11.95/8.86 | +34.87 | <0.0001 | 50/40 |
|  |  |  |  | 3 | 12.30/9.51 | +29.33 | <0.0001 | 50/67 |
| 5h | Length | N2, Mated, Chip | HK/Live OP50 | 1 | 1.25/1.46 | -14.38 | <0.0001 | 20/19 |
|  |  |  |  | 2 | 1.29/1.48 | -12.83 | <0.0001 | 25/25 |
|  |  |  |  | 3 | 1.31/1.49 | -12.08 | <0.0001 | 30/33 |
| 5i | Width | N2, Mated, Chip | HK/Live OP50 | 1 | 61.47/67.76 | -9.28 | 0.0002 | 20/20 |
|  |  |  |  | 2 | 62.38/70.84 | -11.94 | <0.0001 | 25/25 |
|  |  |  |  | 3 | 61.84/66.59 | -7.13 | 0.0001 | 30/33 |
| 5j | LS | *glp-1*, Mated, Chip | HK/Live OP50 | 1 | 9.69/9.02 | +7.42 | 0.3424, n.s. | 50/40 |
|  |  |  |  | 2 | 9.74/9.39 | +3.72 | 0.4001, n.s. | 50/50 |
| 5k | Length | *glp-1*, Mated, Chip | HK/Live OP50 | 1 | 1.46/1.51 | -3.31 | 0.1840, n.s. | 24/25 |
| 6a | RS | *eat-2*, Mated, Chip | HK/Live OP50 | 1 | 14.46/13.54 | +6.79 | 0.0418 | 50/50 |
| 6c | LS | *skn-1*, Mated, Chip | HK/Live OP50 | 1 | 13.16/10.97 | +19.96 | <0.0001 | 62/59 |
|  |  |  |  | 2 | 12.60/11.33 | +11.20 | <0.0001 | 50/40 |
| 6d | RS | *skn-1*, Mated, Chip | HK/Live OP50 | 1 | 9.37/7.67 | +22.16 | 0.0005 | 62/59 |
|  |  |  |  | 2 | 10.32/8.23 | +25.39 | 0.0016 | 42/37 |
| 6e | Length | *skn-1*, Mated, Chip | HK/Live OP50 | 1 | 1.37/1.56 | -13.86 | <0.0001 | 19/19 |
|  |  |  |  | 2 | 1.41/1.55 | -9.03 | <0.0001 | 25/25 |
| 6f | LS | *daf-16*, Mated, Chip | HK/Live OP50 | 1 | 11.17/9.92 | +12.60 | 0.0041 | 50/50 |
|  |  |  |  | 2 | 12.48/10.86 | +14.91 | 0.0002 | 50/50 |
| 6g | RS | *daf-16*, Mated, Chip | HK/Live OP50 | 1 | 10.79/8.16 | +32.23 | <0.0001 | 50/50 |
|  |  |  |  | 2 | 12.04/9.24 | +30.30 | 0.0012 | 50/50 |
| 6h | Length | *daf-16*, Mated, Chip | HK/Live OP50 | 1 | 1.22/1.50 | -18.66 | <0.0001 | 21/20 |
| 6i | LS | *sgk-1*, Mated, Chip | HK/Live OP50 | 1 | 18.43/18.05 | +2.10 | 0.7529, n.s. | 40/40 |
|  |  |  |  | 2 | 15.36/15.00 | +2.40 | 0.8382, n.s. | 50/50 |
| 6j | RS | *sgk-1*, Mated, Chip | HK/Live OP50 | 1 | 15.56/15.81 | -1.58 | 0.7146, n.s. | 40/40 |
|  |  |  |  | 2 | 15.76/14.95 | +5.34 | 0.3483, n.s. | 50/50 |
| 6k | Length | *sgk-1*, Mated, Chip | HK/Live OP50 | 1 | 0.91/0.96 | -4.30 | 0.0103 | 20/20 |
| 7a | LS | Mated, Live OP50, Chip | *sgk-1*/N2 | 1 | 18.05/12.83 | +40.68 | <0.0001 | 40/50 |
|  |  |  | *daf-16*/N2 | 1 | 10.86/12.83 | -15.35 | <0.0001 | 50/50 |
|  |  |  | *skn-1*/N2 | 1 | 10.61/12.83 | -17.30 | <0.0001 | 70/50 |
|  |  |  |  | 2 | 11.33/13.44 | -15.69 | <0.0001 | 40/40 |
| 7b | RS | Mated, Live OP50, Chip | *sgk-1*/N2 | 1 | 15.81/8.53 | +85.34 | <0.0001 | 40/50 |
|  |  |  | *daf-16*/N2 | 1 | 9.24/8.53 | +8.32 | 0.0530, n.s. | 50/50 |
|  |  |  | *skn-1*/N2 | 1 | 6.91/8.53 | -18.99 | 0.0078 | 70/50 |
| 7f | RS/LS | Unmated, Live OP50, Chip | *daf-16*/N2 | NA | 0.54/0.41 | +31.70 | <0.0001 | 41/147 |
|  |  |  | *skn-1*/N2 | NA | 0.48/0.41 | +17.07 | 0.0005 | 28/147 |
|  |  |  | *sgk-1*/N2 | NA | 0.77/0.41 | +87.80 | <0.0001 | 43/147 |
|  |  | Mated, Live OP50, Chip | *sgk-1*/N2 | NA | 0.78/0.66 | +18.18 | <0.0001 | 26/126 |
|  |  | *sgk-1*, Live OP50, Chip | Mated/Unmated | NA | 0.78/0.77 | +1.29 | 0.7899, n.s. | 26/43 |
| 7g | LS | N2, Live OP50, Chip | Mated/Unmated | 1 | 13.08/14.35 | -8.85 | 0.0006 | 67/66 |
|  |  |  |  | 2 | 11.77/13.04 | -9.22 | <0.0001 | 100/100 |
|  |  | *sgk-1*, Live OP50, Chip | Mated/Unmated | 1 | 15.01/14.09 | +6.5 | 0.0003 | 50/50 |
| S.1e | RS | N2, Mated, Live OP50, Chip | repB/repA | NA | 8.53/8.27 | +3.14 | 0.2334, n.s. | 50/100 |
|  |  |  | repC/repA | NA | 7.93/8.27 | -4.11 | 0.2009, n.s. | 99/100 |
|  |  |  | repD/repA | NA | 8.86/8.27 | +7.13 | 0.0610, n.s. | 40/100 |
| S.1f | Length | N2, Live OP50, Chip | Mated/Unmated | 1 | 1.47/1.50 | -2.00 | 0.3953, n.s. | 20/20 |
|  |  |  |  | 2 | 1.48/1.45 | +2.06 | 0.2125, n.s. | 25/25 |
|  |  |  |  | 3 | 1.49/1.51 | -1.32 | 0.5625, n.s. | 33/30 |
| S.1g | LS | N2, Unmated, HK OP50 | NMN/Control | 1 | 11.74/11.74 | +0.02 | 0.3482, n.s. | 100/100 |
| S.1h | RS | N2, Mated, HK OP50 | NMN/Control | 1 | 6.4/6.27 | +1.95 | 0.7553, n.s. | 100/100 |
|  |  |  |  | 2 | 8.14/7.27 | +11.96 | 0.9022, n.s. | 100/100 |
| S.1i | LS | N2, Unmated, HK OP50 | NAC/Control | 1 | 12.72/11.74 | +8.34 | 0.0148 | 100/100 |
| S.1j | RS | N2, Mated, HK OP50 | NAC/Control | 1 | 6.81/6.27 | +8.49 | 0.1998, n.s. | 100/100 |
|  |  |  |  | 2 | 6.24/6.10 | +2.29 | 0.508, n.s. | 85/77 |
| S.3a | RS | N2, Unmated, Chip | HK/Live OP50 | 1 | 5.70/5.39 | +5.75 | 0.0037 | 80/80 |
|  |  |  |  | 2 | 6.23/5.87 | +6.13 | 0.0112 | 50/40 |
| S.3b | Length | N2, Unmated, Chip | HK/Live OP50 | 1 | 1.26/1.50 | -16 | <0.0001 | 21/20 |
|  |  |  |  | 2 | 1.28/1.45 | -11.72 | <0.0001 | 25/25 |
|  |  |  |  | 3 | 1.31/1.51 | -13.24 | <0.0001 | 30/31 |
| S.3c | Width | N2, Unmated, Chip | HK/Live OP50 | 1 | 61.12/71.23 | -14.19 | <0.0001 | 21/20 |
|  |  |  |  | 2 | 62.77/67.44 | -6.92 | 0.0005 | 25/25 |
|  |  |  |  | 3 | 58.46/67.73 | -13.68 | <0.0001 | 30/31 |
| S.3d | Length | Unmated, Live Op50, Chip | *sgk-1*/N2 | 1 | 0.97/1.50 | -35.33 | <0.0001 | 25/20 |

**Table B.**

| **Fig.** | **Test Condition** | **Comparison** | **Experiments** | **n** | **Mean** | **% Change** | **P-value** |
| --- | --- | --- | --- | --- | --- | --- | --- |
| 3a | N2, Mated, Live OP50 | Chip/Plate | 11/7 | 852/519 | 10.86/78.69 | -86.19 | <0.0001 |
| 3b | N2, Unmated, Live OP50 | Chip/Plate | 1 | 100/89 | NA | NA | <0.0001 |
|  | *daf-2*, Unmated, Live OP50 | Chip/Plate | 1 | 196/106 | NA | NA | <0.0001 |
| 5b | N2, Mated, HK OP50 | Chip/Plate | 1 | 89/100 | NA | NA | <0.0001 |
| 6b | *eat-2*, Mated, Chip | HK/Live OP50 | 1 | 50/50 | NA | NA | 0.0125 |

**Table C.**

| **Fig.** | **Correlation** | **Panel Condition** | **Test Condition** | **n** | **Experiments** | **r-value** | **p-value** |
| --- | --- | --- | --- | --- | --- | --- | --- |
| 4a | LS-Length | Unmated, Live OP50, Chip | N2 | 132 | 3 | +0.01 | 0.9313, n.s. |
|  |  |  | *glp-1* | 53 | 2 | -0.33 | 0.0165 |
|  |  |  | *daf-2* | 37 | 1 | +0.38 | 0.0224 |
| 4b | LS-Width | Unmated, Live OP50, Chip | N2 | 132 | 3 | -0.18 | 0.0447 |
|  |  |  | *glp-1* | 53 | 2 | -0.29 | 0.0372 |
|  |  |  | *daf-2* | 37 | 1 | -0.01 | 0.9424, n.s. |
| 4c | RS-Length | N2, Live OP50, Chip | Unmated | 132 | 3 | +0.03 | 0.7241, n.s. |
|  |  |  | Mated | 60 | 2 | -0.29 | 0.0246 |
| 4d | LS-RS | N2, Live OP50, Chip | Unmated | 147 | 3 | +0.02 | 0.7596, n.s. |
|  |  |  | Mated | 126 | 3 | +0.09 | 0.3152, n.s. |
| 4e | LS-PP | N2, Live OP50, Chip | Unmated | 34 | 1 | +0.363 | 0.0346 |
|  |  |  | Mated | 34 | 1 | +0.149 | 0.4003, n.s. |
| 4f | RS-PP | N2, Live OP50, Chip | Unmated | 50 | 1 | +0.543 | <0.0001 |
|  |  |  | Mated | 49 | 1 | +0.917 | <0.0001 |
| 5d | LS-RS | N2, Unmated, Chip | Live OP50 | 147 | 3 | +0.02 | 0.7596, n.s. |
|  |  |  | HK OP50 | 168 | 3 | -0.06 | 0.4406, n.s. |
| 5g | LS-RS | N2, Mated, Chip | Live OP50 | 126 | 3 | +0.09 | 0.3152, n.s. |
|  |  |  | HK OP50 | 132 | 3 | +0.25 | 0.0045 |
| 7c | LS-RS | Unmated, Live OP50, Chip | *sgk-1* | 43 | 1 | +0.49 | 0.0008 |
|  |  |  | N2 | 147 | 3 | +0.02 | 0.7596, n.s. |
|  |  |  | *skn-1* | 28 | 1 | -0.06 | 0.7432, n.s. |
|  |  |  | *daf-16* | 41 | 1 | -0.26 | 0.0901, n.s. |
| 7d | LS-RS | Mated, Live OP50, Chip | *sgk-1* | 26 | 2 | +0.64 | 0.0004 |
|  |  |  | N2 | 126 | 3 | +0.09 | 0.3152, n.s. |
|  |  |  | *skn-1* | 104 | 3 | +0.50 | <0.0001 |
|  |  |  | *daf-16* | 48 | 2 | +0.55 | <0.0001 |

**Figure S1:** (a) An incubation arena can house multiple worms in lifespan assay. (b) *Ce*Lab chip. (c) *Ce*Lab control center next to a dissecting scope for manual scoring. (d) Cross-section of *Ce*Lab's multi-layer design. (e) *Ce*Lab‘s reproductive span data are highly reproducible, (repA, n=100; repB, n=50; repC, n=99; repD, n=40). (f) Culturing in liquid abrogates mating-induced water loss-related shrinking, (unmated, n=20; Mated, n=20). (g, h) lifespan and reproductive span of animals treated with NAC (control, NAC, n=100 in both g and h) and (i, j) NMN (control, NMN, n=100 in both i and j). Kaplan-Meier survival tests. Two-tailed t-tests. ns not significant. Box plots show minimum, 25th percentile, median, 75th percentile, maximum.

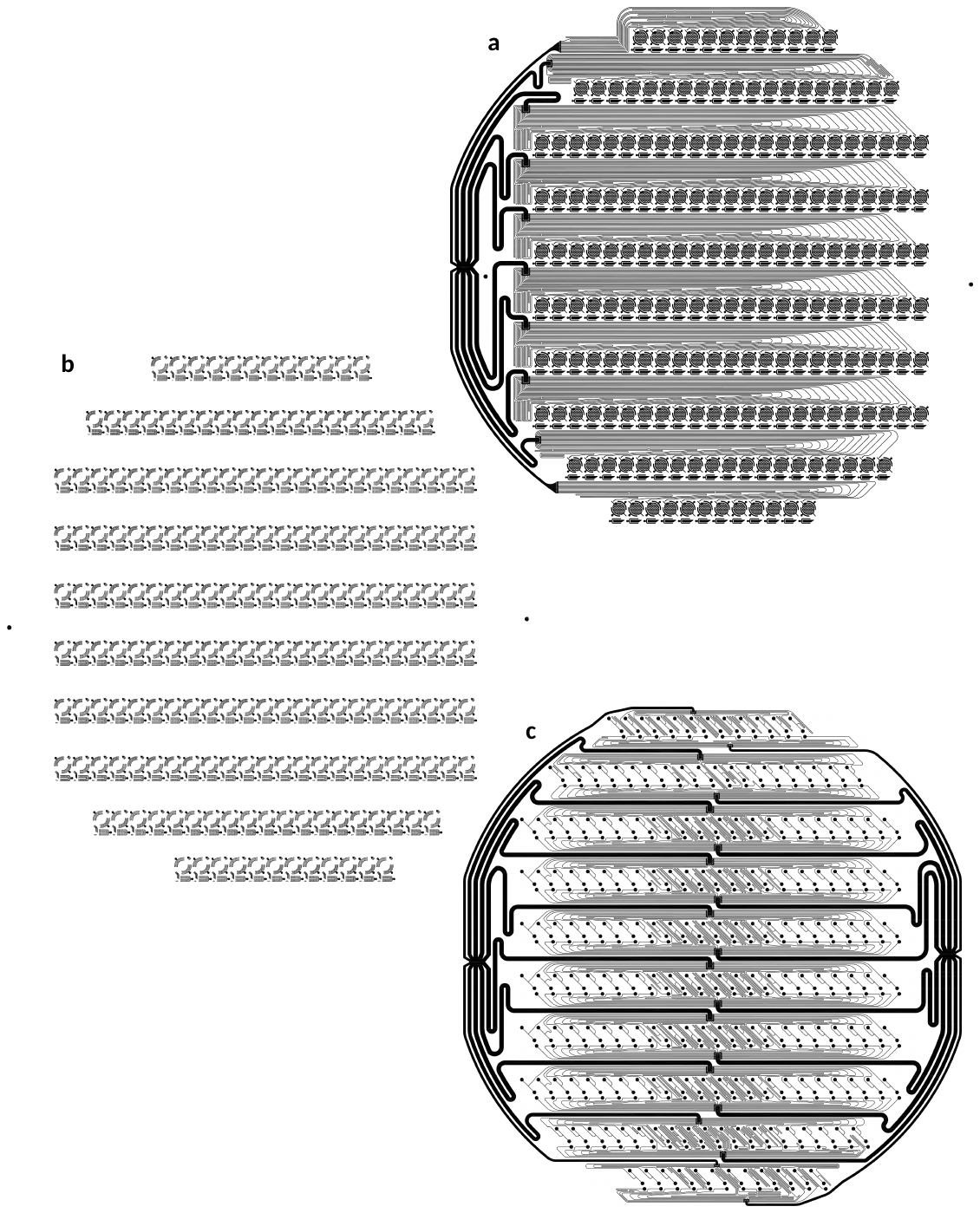

**Figure S2.** Mask designs for Layers (a) #3, (b) #4, and (c) #5.

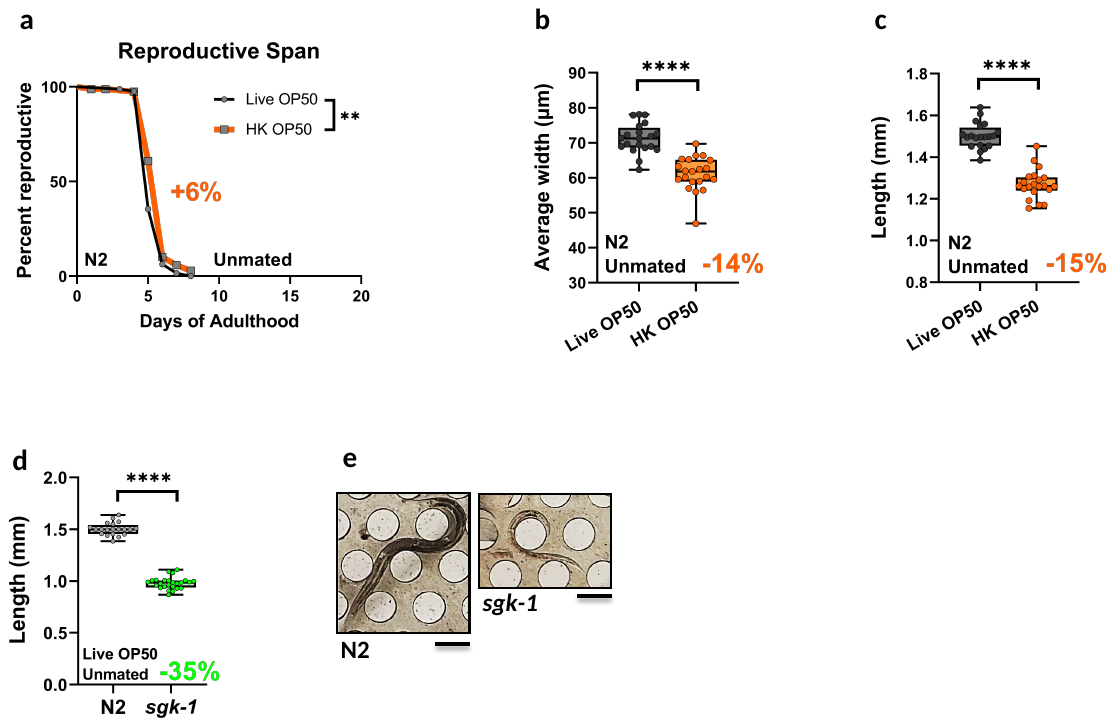

**Figure S3:** (a) Reproductive span of unmated wild-type animals is slightly increased on HK diet, (Live, n=80; HK, n=80). Unmated wild-type worms on HK bacteria are significantly shorter, (Live, n=20; HK, n=21) and (i) thinner, (Live, n=20; HK, n=21). (d, e) *sgk-1* mutants are significantly smaller than wild-type animals, (N2, n=20; *sgk-1*, n=25). Scale bar, 200 µm. Kaplan-Meier survival tests. Two-tailed t-tests. **p < 0.01, ****p < 0.0001. Box plots show minimum, 25th percentile, median, 75th percentile, maximum.

**Video Legends**

Video S1. *Ce*Lab operation workflow. (a) Loading animals. (b, c) Progeny and waste removal. (d) Feeding. (e) Scoring.

Video S2. *Ce*Lab allows analysis of movement, which can be used to carry out motility assays.
